# Nutritional vitamin B12 regulates RAS/MAPK-mediated cell fate decisions through the one-carbon metabolism

**DOI:** 10.1101/2023.10.12.562015

**Authors:** Ana Cristina Laranjeira, Simon Berger, Tea Kohlbrenner, Nadja R. Greter, Alex Hajnal

**Affiliations:** Institute of Molecular Life Sciences, University of Zurich, Winterthurerstrasse 190, Zurich, Switzerland; Institute for Chemical and Bioengineering, ETH Zurich, Vladimir-Prelog-Weg 1-5/10, Zurich, Switzerland; PhD program in Molecular Life Science

**Keywords:** RAS/MAPK, one-carbon metabolism, vitamin B12, methionine cycle, choline, H3K4me3, fatty acids, cell fate decision

## Abstract

Vitamin B12 is an essential nutritional co-factor for the folate and methionine cycles, which together constitute the one-carbon metabolism. Here, we show that dietary uptake of vitamin B12 modulates cell fate decisions controlled by the conserved RAS/MAPK signaling pathway in *C. elegans*. A bacterial diet rich in vitamin B12 increases vulval induction, germ cell apoptosis and oocyte differentiation. These effects are mediated by different one-carbon metabolites in a tissue-specific manner. Vitamin B12 enhances via the choline/phosphatidylcholine metabolism vulval induction by down-regulating fat biosynthesis genes and increasing H3K4 tri-methylation, which results in increased expression of RAS/MAPK target genes. Furthermore, the nucleotide metabolism and H3K4 tri-methylation positively regulate germ cell apoptosis and oocyte production. Using mammalian cells carrying different activated KRAS and BRAF alleles, we show that the effects of methionine on RAS/MAPK-regulated phenotype are conserved in mammals. Our findings suggest that the vitamin B12-dependent one-carbon metabolism is a limiting factor for diverse RAS/MAPK-induced cellular responses.

## Introduction

The cell metabolism regulates physiological processes and cancer cell growth by controlling catabolic and anabolic reactions and transcriptional activity (Bose et al., 2020; Hanahan and Weinberg, 2011). Diet influences metabolism directly and, as a result, cells must constantly adapt to variations in diet. One example is vitamin B12 (cobalamin), an essential micronutrient that requires dietary intake. Vitamin B12 is an important metabolic co-factor in two metabolic pathways, the mitochondrial propionate breakdown pathway and the one-carbon metabolism formed by the methionine and folate cycles (**Figure 1A**). In the canonical propionate breakdown pathway, vitamin B12 acts as a co-factor for methyl malonyl-CoA mutase (MUT, MMCM-1 in *C. elegans*) (Bito and Watanabe, 2016; Green et al., 2017; Watson et al., 2014, 2016), and in the methionine cycle, it serves as a co-factor for methionine synthase (MS, METR-1 in *C. elegans*) (**Figure 1A**) (Froese et al., 2019; Green et al., 2017).

**Figure 1.**
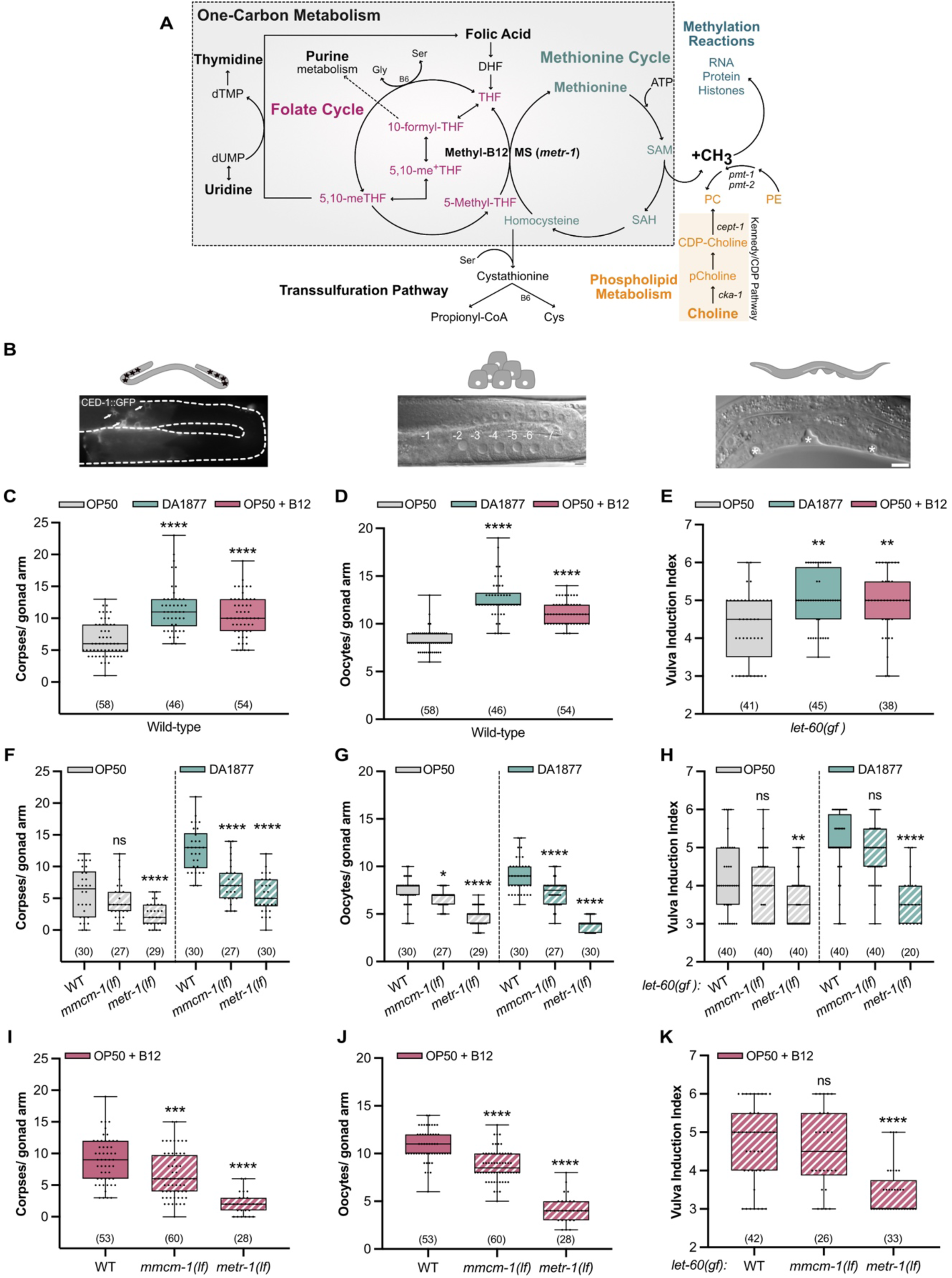
The DA1877 diet promotes VPC and germ cell differentiation through the vitamin B12 metabolism. (A) Schematic representation of the one-carbon metabolism. (B) Illustrations of the observed phenotypes: germ cell apoptosis (corpses), oocyte differentiation or vulval induction; scale bar: 10 µm. (C-D) Number of corpses (C) or oocytes (D) in wild-type animals. (E) Vulval induction index of *let-60(gf)* animals. (C-E) Animals were fed with OP50 (gray), DA1877 (blue) or OP50 supplemented with 64 nM B12 (pink). (F-H) Number of corpses (F), oocytes (G) and vulval induction index (H) for indicated genotypes fed either with OP50 (gray) or DA1877 (blue). (I-K) Number of corpses (I), oocytes (J) and vulval induction index (K) for indicated genotypes fed with OP50 and supplemented with 64 nM B12. Dots (in C, D, F, G, I, J) represent individual corpses/oocytes or animals (in E, H, K) from two independent biological replicates; number of animals in brackets; data represent median ± min and max. For statistical analysis see STAR methods. **** <0.0001; ***<0.0005; **<0.005; ns, non-significant.

The one-carbon metabolism transfers single carbon units necessary for nucleotide biosynthesis through the folate cycle and produces S-adenosyl-methionine (SAM), the main methyl donor in cells. In the methionine cycle, homocysteine is methylated by MS to form methionine, which is then converted into SAM (Ducker and Rabinowitz, 2017; Newman and Maddocks, 2017; Ye et al., 2017). SAM is required for phosphatidylcholine (PC) synthesis (**Figure 1A**) (Ducker and Rabinowitz, 2017; Froese et al., 2019; Newman and Maddocks, 2017). PC is a polyunsaturated fatty acid that can be produced via the Kennedy/CDP-choline pathway using choline as a precursor, or by the sequential methylation pathway, which depends on SAM-methyl groups (**Figure 1A**) (Brendza et al., 2007; Cui and Houweling, 2002; Lochnit and Geyer, 2003). In addition, SAM provides the methyl donors necessary for DNA, RNA, histone and protein methylation. Histone methylation, especially histone H3 tri-methylation at lysine 4 (H3K4me3), which is associated with active gene expression, is particularly sensitive to SAM levels (Ding et al., 2015; Mentch et al., 2015; Shyh-Chang et al., 2013).

In this study, we have used the nematode *C. elegans* to investigate the nutritional impact of a bacterial diet on development. *E. coli* OP50 is the commonly used bacterial diet, which is low in vitamin B12. *Comamonas aquosa* DA1877, on the other hand, is a vitamin B12-rich diet (Watson et al., 2014). A DA1877 diet affects different aspects of *C. elegans* behavior by modulating gene expression, reproduction and longevity (MacNeil et al., 2013; Watson et al., 2013). The vitamin B12 metabolism also regulates H3K4 methylation (Ding et al., 2015), affecting *C. elegans* fertility and lipid accumulation, which is linked to longevity (Ding et al., 2018; Greer et al., 2010; Xiao et al., 2011).

Here, we focused on how the vitamin B12-dependent one-carbon metabolism modulates cell fate decisions mediated by the conserved RAS/Mitogen-activated protein kinase (RAS/MAPK) pathway. The RAS/MAPK pathway is highly conserved between *C. elegans* and mammals, and it is a prevalent oncogenic signaling pathway in humans, being hyper-activated in around one-third of all tumors. Gain-of-function (*gf*) mutations in RAS genes (mostly in KRAS) occur in 30% of all cancers, and mutations in BRAF (mostly the V600E substitution) are found in 8% of all human cancers (Guo et al., 2020; Liu et al., 2018; Schubbert et al., 2007). Gain-of-function mutations in *let-60*, the single *C. elegans ras* ortholog, hyperactivate RAS/MAPK signaling and cause enhanced germ cell death, oocyte over-production and excess vulval induction (Cha et al., 2012; Church et al., 1995; Han et al., 1990; Lee et al., 2007; Sternberg and Han, 1998).

During larval development, the growth factor LIN-3 (an EGF-like ligand) is secreted by the gonadal anchor cell (AC). LIN-3 activates RAS/MAPK signaling via the LET-23 EGF receptor in the vulval precursor cells (VPCs) to induce their differentiation (Greenwald, 1997; Gupta et al., 2012; Schindler and Sherwood, 2013). P6.p, the VPC closest to the AC, receives the highest concentration of LIN-3 and adopts the primary (1°) vulval cell fate, while P5.p and P7.p receive less LIN-3 signal and adopt the secondary (2°) cell fate in response to a lateral LIN-12 Notch signal. These three induced VPCs then start to proliferate and differentiate into 22 vulval cells forming the vulva (Greenwald, 1997; Gupta et al., 2012; Schindler and Sherwood, 2013).

During adulthood, RAS/MAPK regulates germ cell progression and death in the gonads. Hermaphrodites have two symmetric, U-shaped tubular gonad arms that are connected to a common uterus. In the distal region of each gonad arm, mitotic stem cells constantly proliferate to supply new germ cells. Germ cells in the late pachytene region, near the gonad loop, activate the RAS/MAPK pathway, which is required for the pachytene to diplotene transition, cell membrane integrity, chromosomal synapsis, apoptosis and oocyte differentiation (Arur et al., 2011; Church et al., 1995; Das et al., 2020; Lee et al., 2007). Around half of the germ cells enter the proximal gonad arm and begin to differentiate into oocytes, while the remaining germ cells undergo apoptosis. Germ cell apoptosis is a physiological and stochastic process that eliminates half of all germ cells to maintain gonad homeostasis (Gumienny et al., 1999; Wang and Yang, 2016). Dying germ cells activate the core apoptotic machinery triggering the CED-3 caspase (Gumienny et al., 1999; Wang and Yang, 2016) and are engulfed by the sheath cells of the somatic gonad, which express the CED-1 receptor mediating their recognition (Zhou et al., 2001).

By examining three RAS/MAPK-mediated phenotypes, vulval development, germ cell death and oocyte differentiation, we show that the activity of the vitamin B12-dependent one-carbon metabolism is a limiting factor for RAS/MAPK-induced cell fates. Switching animals to a vitamin B12-rich DA1877 diet enhances all phenotypes caused by hyperactive RAS/MAPK signaling via the methionine synthetase METR-1. The one-carbon metabolism affects the different RAS/MAPK-mediated cell fate decisions through distinct, tissue-specific mechanisms. The folate cycle promotes germ cell differentiation and death through nucleotide biosynthesis, while the methionine cycle affects vulval induction by repressing fatty acid synthesis. In addition, the methionine cycle regulates the PC metabolism, which in turn increases global H3K4me3 levels affecting all tissues. Finally, using mammalian cells carrying different activating KRAS mutations, we show that the methionine-dependency of RAS/MAPK-induced phenotypes is conserved.

## Results

### A vitamin B12-rich diet enhances germ cell apoptosis, oocyte differentiation and vulval induction

To investigate how the vitamin B12 metabolism affects RAS/MAPK-controlled cell fate decisions, we fed animals with the DA1877 diet, rich in vitamin B12, and quantified germ cell death, oocyte and VPC differentiation (**Figure 1B**). Using a CED-1::GFP reporter to label engulfed apoptotic cells (Derry et al., 2001), we found that wild-type worms fed with the DA1877 diet had a 2 to 3-fold increase in apoptotic germ cells, and more oocytes in the proximal gonad arm (**Figures 1C** and **1D**). Supplementing vitamin B12 to animals grown on OP50 bacteria likewise increased germ cell apoptosis and oocyte numbers (**Figures 1C** and **1D**).

Since DA1877-fed worms reach adulthood on average 12 hours earlier (**Figure S1A**) (Watson et al., 2013), we tested whether these effects were due to changes in developmental timing. Sixty-two hours after L1 arrest, DA1877-fed animals were approximately at the same developmental stage as animals fed on OP50 for 72 hours. However, DA1877-fed worms had more apoptotic corpses and oocytes than OP50-fed animals at the matching developmental stages (i.e., 62 vs. 72h, **Figures S1B** and **S1C**), suggesting that the DA1877 effects are independent of developmental time.

Germ cell apoptosis can either be physiological or a response to DNA damage (Gartner et al., 2000; Gumienny et al., 1999). To distinguish between these two possibilities, we used *cep-1* loss-of-function (*lf*) mutants, which do not induce DNA damage-induced apoptosis (Derry et al., 2001). The effect of the DA1877 diet on germ cell death was still present in *cep-1(lf)* mutants (**Figure S1D**). We also examined *pch-2(lf)* mutants (Kohlbrenner et al., 2023) to exclude that the enhanced germ cell apoptosis in DA1877-fed animals was due to defects in synaptonemal complex assembly (**Figure S1E**). Thus, the DA1877 diet enhances physiological germ cell death. In contrast, a *ced-3(lf)* mutation completely suppressed the effect of the DA1877 diet, indicating that the canonical apoptotic caspase pathway is activated by the diet (**Figure S1F**).

To study the effect of the DA1877 diet on VPC induction during vulval development, we used a *let-60 ras* gain-of-function mutation (*let-60(n1046)*, abbreviated *let-60(gf*)), as a sensitized genetic background. *let-60(gf*) animals grown on OP50 contained on average 4.3±0.9 (SD) induced VPCs per animal (vulval induction index). *let-60(gf*) larvae fed with DA1877 or supplemented with vitamin B12 showed an increase in vulval induction to 5.0±0.7 and 5.0±0.8, respectively (**Figure 1E**). Wild-type larvae always had three induced VPCs irrespective of the bacterial diet.

To test if the effect of the DA1877 diet was caused by bacterially produced B12, we used the B12 synthesis-deficient DA1877 mutants *ΔcbiA/cobB* and *ΔcbiB* (Watson et al., 2014). Germ cell apoptosis in animals fed with DA1877 *ΔcbiA/cobB* or *ΔcbiB* mutants was comparable to OP50-fed animals, indicating that the increase in dietary B12 is the main cause for the enhanced apoptosis (**Figure S1G**). To determine, which metabolic pathway (i.e., the propionate breakdown or the one-carbon metabolism) mediates the effects of the DA1877 diet or B12 supplementation, we examined *metr-1(lf)* and *mmcm-1(lf)* mutants. Both mutations suppressed the effect of the DA1877 diet and vitamin B12 on germ cell apoptosis and oocyte differentiation, though the suppression by *mmcm-1(lf)* was weaker (**Figures 1F, 1G, 1I and 1J**). Only *metr-1(lf)* mutants showed a reduction in VPC induction in animals fed with the DA1877 diet or supplemented with vitamin B12 (**Figures 1H** and **1K**). To determine if the effects of vitamin B12 on germ cell differentiation are cell-autonomous, we expressed *metr-1* and *mmcm-1* in the soma using extra-chromosomal arrays, which are silenced in the germline. The somatic expression of *metr-1* rescued the reduced vulval induction in *let-60(gf)* double mutants (**Figure S1H**), but the somatic expression of *metr-1* or *mmcm-1* did not affect germ cell death on the DA1877 diet (**Figure S1I**).

Lastly, we examined duct cell specification during larval development as a fourth phenotype controlled by RAS/MAPK signaling (Sternberg and Han, 1998). Eighty-one percent of *let-60(gf)* animals fed with OP50 showed a duplication of the duct cell, whereas 98% of the animals fed with the DA1877 diet had two duct cells **(Figures S1J** and **S1J’**).

In conclusion, the vitamin B12-dependent one-carbon metabolism is the main factor enhancing multiple RAS/MAPK-induced cell fates on the DA1877 diet.

### A vitamin B12-rich diet increases germ cell number and accelerates pachytene progression

Given that the DA1877 diet affected both germ cell death and oocyte differentiation, we tried to decouple the two phenotypes. Using apoptosis-deficient *ced-3(lf)* mutants, we found that the increase in oocyte number in DA1877-fed worms was independent of apoptosis (**Figure 2A**). However, we were not able to determine if the reverse was also true.

**Figure 2.**
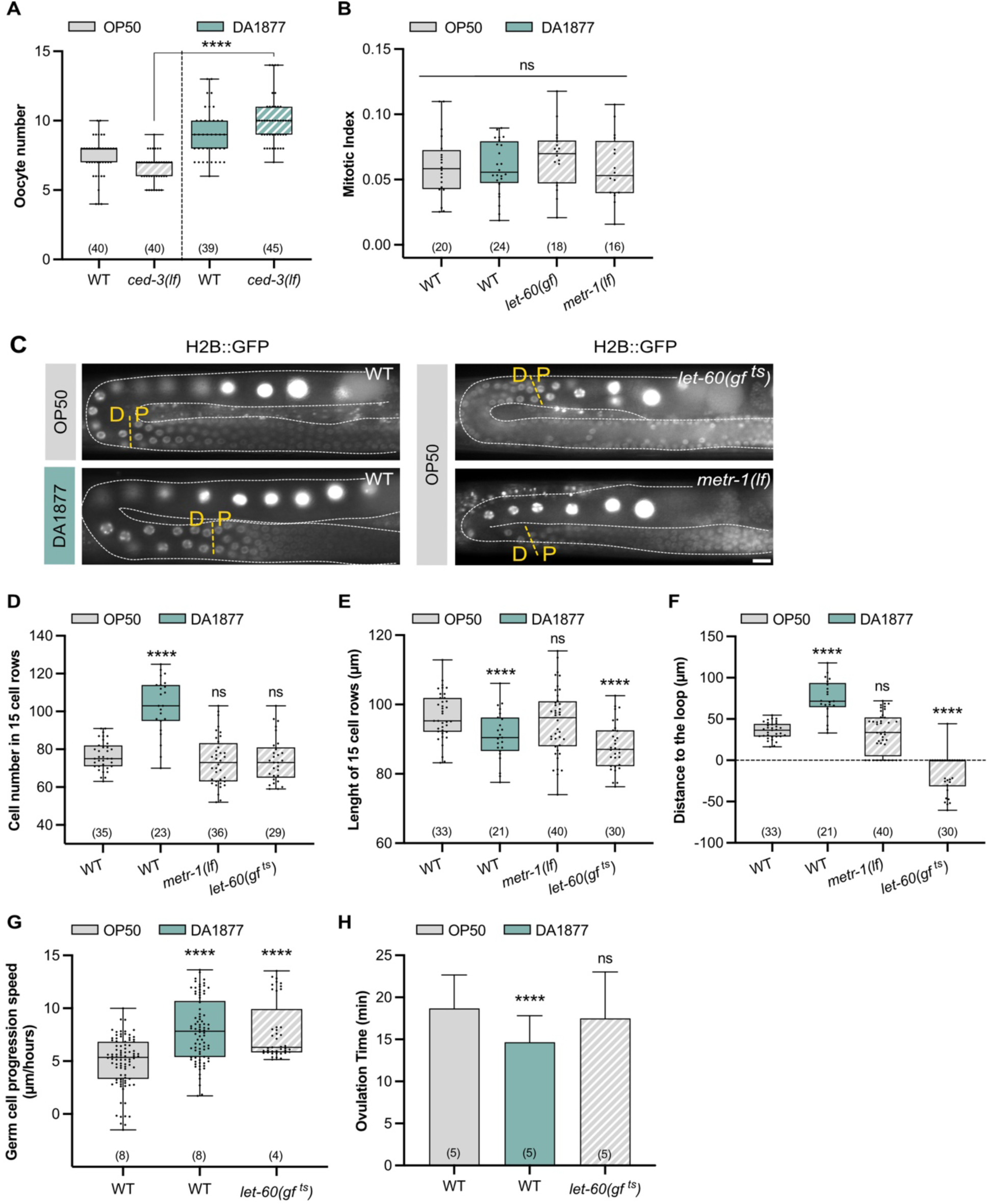
The DA1877 diet changes germ cell number and dynamics. (A) Oocyte number for indicated genotypes; dots represent individual oocytes. (B) Mitotic index for indicated genotypes; dots represent individual animals. (C) Fluorescence images showing the pachytene-to-diplotene (P-D) transition (yellow dashed lines) for indicated genotypes; germ cell nuclei are marked with H2B::GFP; gonads are outlined by white dashed lines; scale bar: 10 µm. (D) Number of cells inside an area starting from the P-D and ending 15 cell-rows distally of indicated genotypes; P-D was determined based on the nuclear marker shown in (C); in *let-60(gf)* animals the loop region was used as the starting point. (E) Length of 15-cell rows for indicated genotypes. (F) Distance from the P-D to the loop region for indicated genotypes. (D-F) dots represent individual animals. (G) Speed of germ cell progression through the pachytene region for indicated genotypes; for individual tracking see Figure S2; dots represent the speed of individual cells. (H) Ovulation time for indicated genotypes; dots represent individual oocytes; bars represent mean ± SD. (A-H) Animals were fed with OP50 (gray) or DA1877 (blue). Number of animals in brackets; data represent median ± min and max. For statistical analysis see STAR methods. **** <0.0001; ns, non-significant.

The mitotic proliferation of germline stem cells in the distal gonad region depends on RAS/MAPK signaling in the soma (Robinson-Thiewes et al., 2021). We therefore tested if the DA1877 diet up-regulates germ cell proliferation, which would increase germ cell number and potentially apoptosis. The mitotic index (i.e. the fraction of M-phase germ cells in the mitotic region) in adult animals was not changed by the DA1877 diet, by somatic hyper-activation of RAS/MAPK, or by loss of *metr-1* (**Figure 2B**). However, DA1877-fed worms contained a longer mitotic zone with more mitotic cells and an increased absolute number of cells in M-phase (**Figure S2A**), which is consistent with the overall increase in gonad size observed in DA1877-fed animals. By contrast, *let-60(gf)* and *metr-1(lf)* mutants contained a shorter mitotic zone (**Figure S2A**). This suggests that the DA1877 diet may affect the germline by expanding the pool of mitotic cells earlier during larval development without increasing the mitotic rate during adulthood. Thus, accelerated germ cell proliferation is unlikely to be responsible for the DA1877-induced increase in apoptotic corpse and oocyte numbers.

We next focused on the pachytene region, where RAS/MAPK activation regulates both germ cell apoptosis and pachytene exit (Church et al., 1995; Gumienny et al., 1999; Lee et al., 2007). We used an H2B::GFP marker to label the germ cell nuclei and determine the border between the pachytene and diplotene zones (P-D) (**Figure 2C**). We counted the average number of pachytene cells over 15-cell rows distal to the P-D border. DA1877-fed animals contained more pachytene germ cells (**Figure 2D**), consistent with an enlarged gonad. To hyper-activate RAS/MAPK signaling in the germline, we examined temperature-sensitive *let-60(ga89)* mutants, abbreviated *let-60(gf ^ts^)*, grown for 4 hours at 25°C (Eisenmann and Kim, 1997). The *let-60(gf ^ts^)* and *metr-1(lf)* mutations did not significantly alter germ cell number (**Figure 2D**). DA1877-fed worm and *let-60(gf ^ts^)* mutants also showed an increased density of pachytene cells (i.e., shorter 15-cell rows), but cell density was not changed in *metr-1(lf)* mutants (**Figure 2E**). Thus, neither cell number nor density seems to correlate with germ cell apoptosis or oocyte differentiation. However, DA1877-fed worms showed a distal shift of the P-D border before the loop, whereas *let-60(gf ^ts^)* mutants showed a proximal shift after the loop (**Figures 2C** and 2**F**).

The distal shift of the P-D border in DA1877-fed worms might be due to a change in germline length, a shift of the borders between the different meiotic zones, or altered dynamics of the germ cells progressing through the gonads. Immunohistochemical staining of dissected gonads with markers for the different meiotic zones indicated that the boundary between the transition zone and early pachytene was shifted distally in DA1877-fed animals (when corrected for different gonad length), but the proximal border of the late pachytene zone was not changed (**Figure S2B**). By contrast, the nuclear morphology visualized with the H2B::GFP marker pointed to a distal shift of the P-D border (**Figure 2C** and **2F)**. In agreement with previous results (Das et al., 2020), *let-60(gf ^ts^)* worms had a shorter mid-pachytene zone and no change in the proximal border of the late pachytene zone, despite the proximal shift of the P-D border determined with the H2B::GFP marker (**Figure S2B** and **Figures 2C** and **2F**). This suggests that the different meiotic zones determined by molecular markers do not always match the meiotic stage of individual germ cells scored by their nuclear morphology.

We therefore hypothesized that the changes in germ cell death and differentiation caused by the DA1877 diet or the *let-60(gf ^ts^)* mutation might be caused by an altered speed, at which individual germ cells move through the different meiotic zones. We tracked individual germ cell nuclei progressing from pachytene to diplotene by live imaging (Berger et al., 2018). Germ cells progressed faster through the pachytene region in DA1877-fed animals and *let-60(gf ^ts^)* mutants (**Figure 2G** and **Figure S3A** and **S3B** for the individual tracking). Moreover, DA1877-fed worms ovulated more frequently (**Figure 2H** and **Figure S3C** for the individual tracking), indicating that the higher oocyte number is not due to reduced ovulation.

In summary, the DA1877 diet or hyper-activation of the RAS/MAPK pathway resulted in a faster progression of germ cells through pachytene, which could explain the increased number of oocytes in the proximal gonads. Whether the accelerated progression through pachytene might also increase the rate of germ cell apoptosis is unclear. It seems more likely that the combination of the changes in gonad morphology and dynamics caused by the DA1877 diet accounts for the increase in germ cell death.

### The one-carbon metabolism enhances RAS/MAPK target gene expression rather than pathway activity

To characterize the interactions between the one-carbon metabolism and RAS/MAPK signaling, we combined the *metr-1(lf)* and *let-60(gf)* mutations. *metr-1(lf)* rescued the *let-60(gf ^ts^)*-dependent increase in germ cell apoptosis and oocyte number in animals fed with OP50, even in germline-specific *metr-1(lf)* mutants rescued in the soma with an extra-chromosomal array (**Figures 3A** and **3B** and **Figures S4A** and **S4B**). Furthermore, the gonads in *let-60(gf ^ts^)* animals grown for 18 hours at 25°C shrank, resulting in a reduced rachis diameter, which was suppressed by *metr-1(lf)* (**Figure 3C**) (Kohlbrenner et al., 2023; Lee et al., 2007). Also, the increased VPC differentiation in *let-60(gf)* mutants fed with OP50 was suppressed by *metr-1(lf)* (**Figures 3D** and **1H**). Together, these results show that inhibiting the methionine cycle reduces the penetrance of phenotypes caused by RAS/MAPK hyper-activation in different tissues. On the other hand, the DA1877 diet enhanced the effects of *let-60(gf ^ts^)* on germ cell apoptosis and oocyte differentiation (**Figures 3E** and **3F**). To examine the influence of the DA1877 diet on reduced RAS/MAPK signaling, we used the temperature-sensitive *mpk-1(ga111 ^ts^)* allele (Lackner and Kim, 1998), rescued in the soma with an extra-chromosomal *mpk-1(+)* array to inhibit RAS/MAPK specifically in the germline (Kohlbrenner et al., 2023). The reduced germ cell apoptosis and oocyte number in germline-specific *mpk-1(lf ^ts^)* mutants were partially suppressed by the DA1877 diet (**Figures 3G** and **3H**). This suggested that the RAS/MAPK pathway and the methionine cycle have synergistic effects, or that the methionine cycle performs a limiting function downstream of the RAS/MAPK pathway.

**Figure 3.**
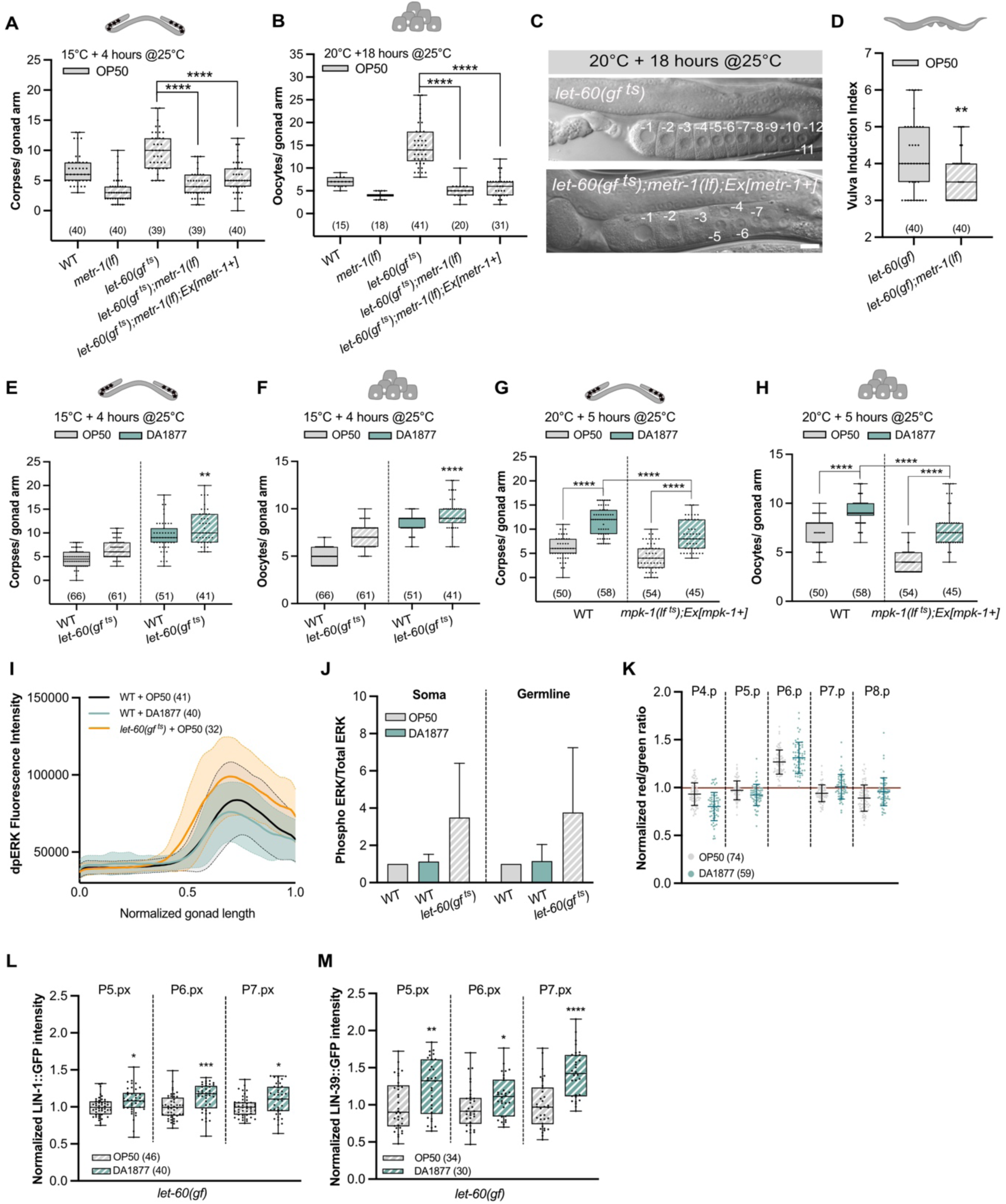
Interaction of the one-carbon metabolism and the RAS/MAPK pathway. (A-B) Number of corpses for indicated genotypes grown at 15°C and transferred to 25°C 4 hours before quantification (A) and oocyte number for indicated genotypes grown at 20°C and transferred to 25°C 18 hours before quantification (B). (C) DIC images of gonads for indicated genotypes grown at 20°C and transferred to 25°C 18 hours before quantification; numbers indicate mature oocytes; scale bar: 10 µm. (D) Vulval induction index for indicated genotypes. (A-D) Animals were fed with OP50. (E-F) Number of corpses (E) and oocytes (F) for indicated genotypes grown at 15°C and transferred to 25°C 4 hours before quantification. (G-H) Number of corpses (G) or oocytes (H) for indicated genotypes grown at 20°C and transferred to 25°C 5 hours before quantification. (E-H) Animals were fed with OP50 (gray) or DA1877 (blue). (I) Summed intensity profile of dpERK staining along the distal-proximal axis of dissected gonads of wild-type fed with OP50 (gray) or DA1877 (blue), and *let-60(gf ^ts^)* fed with OP50 (orange); x-axis represents normalized gonad length, from distal (x=0) to loop (x=1); the full line represents the average intensity ± SD (shaded); data from three independent biological replicates. (J) Western blot quantification of the somatic or germline isoform of phosphorylated and total ERK for indicated genotypes; bars represent mean ± SD normalized to the control (wild-type OP50) from three independent biological and technical replicates; see Figure S3 for the individual Western blots. (K) ERK-nKTR biosensor quantification; in the VPCs normalized red/green ratio in individual VPCs; dots represent different animals from 2 independent biological replicates. (L-M) Normalized intensity of LIN-1::GFP (L) or LIN-39::GFP (M) in the two-cell (Pn.px) stage in 1° and 2° VPCs of *let-60(gf)* animals; data were normalized to the control (OP50); dots represent individual animals from two independent biological replicates. (J-M) Animals were fed with OP50 (gray) or DA1877 (blue). Number of animals is shown in brackets; data represent median ± min and max. For statistical analysis, see STAR methods. **** <0.0001; ***<0.0005; **<0.005; *<0.05.

We next examined if the methionine cycle directly regulates RAS/MAPK pathway activity. Neither immunostaining of dissected gonads nor Western blot analysis of total animal extracts with a di-phospho-ERK-specific antibody showed an increase in MAPK phosphorylation in animals fed with the DA1877 diet (**Figure 3I** and **3J**, see **Figure S4C** for the Western blots used for quantification). Accordingly, *metr-1(lf)* did not cause a decrease in MAPK phosphorylation (**Figures S4D** and **S4E**). Using an ERK biosensor to quantify RAS/MAPK pathway activity in the VPCs (de la Cova et al., 2017), we observed no significant difference in activity between worms fed with the OP50 or DA1877 diet (**Figure 3K**). Together, these results indicate that the DA1877 diet does not increase RAS/MAPK pathway activity but rather the cellular responses to MAPK activation.

We thus tested whether the DA1877 diet changes the expression of RAS/MAPK signaling target genes during vulval development. We used endogenous GFP reporters for *lin-1*, which encodes an ETS-family transcription factor phosphorylated by MPK-1, and *lin-39*, which encodes a *hox* gene regulated by LIN-1 (Dave et al., 1998; Sundaram, 2013). The DA1877 diet increased LIN-1::GFP and LIN-39::GFP expression in the proximal VPCs of *let-60(gf)* larvae (**Figures 3L** and **3M**), but it had no significant effect in wild-type worms (**Figures S4F** and **S4G**).

Thus, the interaction between the methionine cycle and the RAS/MAPK pathway most likely occurs at the level of the target genes. Furthermore, the methionine cycle appears to limit vulval induction only in the context of a hyper-activated RAS/MAPK pathway.

### Tissue-specific effects of one-carbon metabolites on RAS/MAPK-mediated cell fates

To characterize the effects of bacterial diet on the metabolism, we used liquid chromatography mass spectrometry (LC-MS) analysis to measure metabolite levels in animals fed with the OP50 or DA1877 diet. We focused on vitamin B12-related metabolic pathways and found that the DA1877 diet, as expected, increased the levels of one-carbon metabolites (**Figure 4A**, **Table S1** and **Figure S5A**). Particularly, methionine, phosphatidylcholine (PC) and nucleotide levels were increased in DA1877-fed wild-type animals (**Figure S5A**) (Giese et al., 2020). Interestingly, *let-60(gf ^ts^)* mutants fed with the DA1877 diet contained lower levels of one-carbon metabolites than wild-type animals on DA1877 (**Figure 4A** and **S5A**).

**Figure 4.**
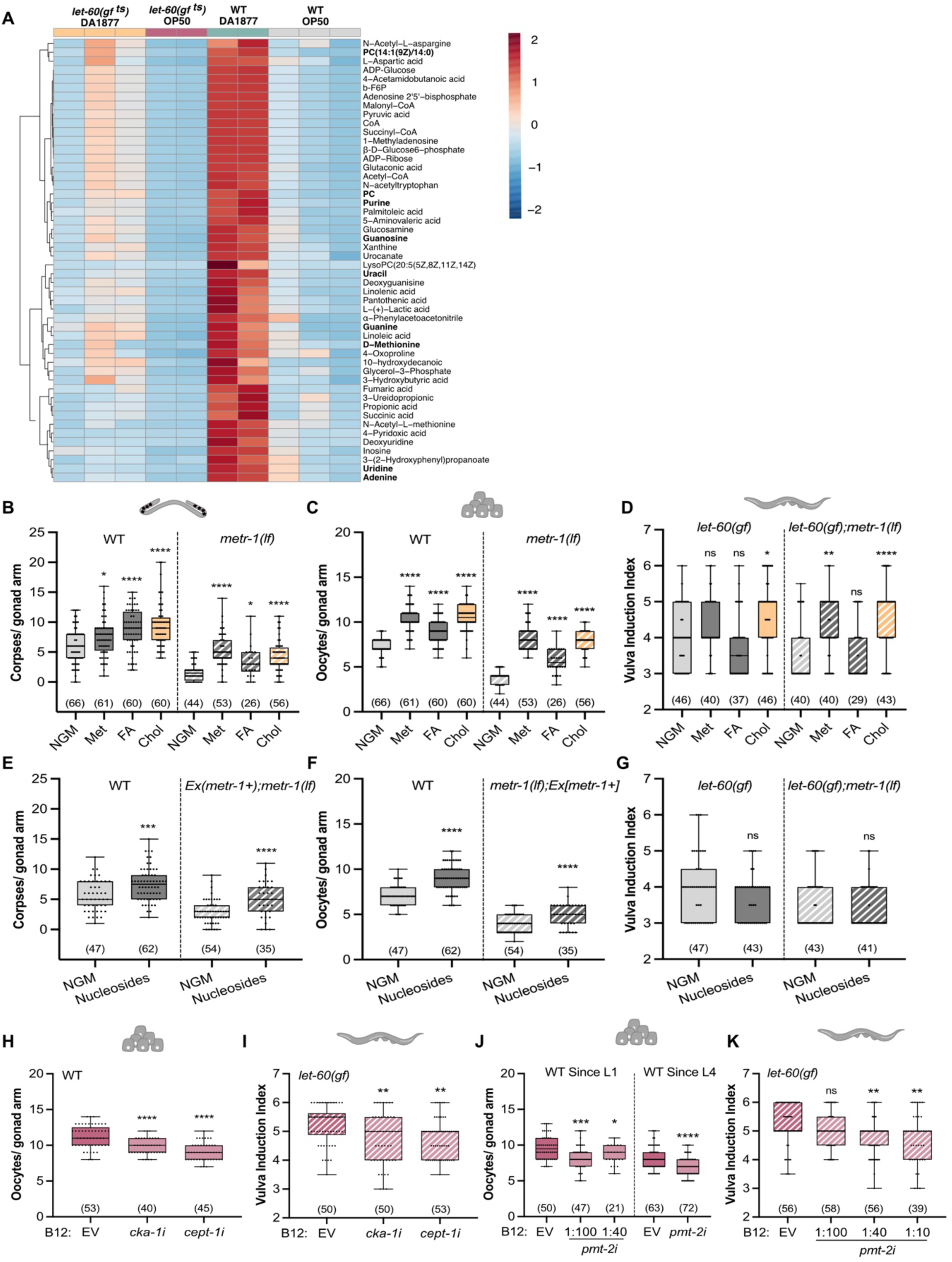
One-carbon metabolites regulate VPC and germline differentiation. (A) Heat map of top 50 changed metabolites of the vitamin B12-dependent metabolism for indicated genotypes; *let-60(gf ^ts^)* animals were transferred to 25°C 4 hours before sample collection; statistical analysis using a one-way ANOVA, considering a minimum p-value of 0.05. (B-G) Number of corpses (B, E), oocytes (C, F) and vulval induction index (D, G) for indicated genotypes fed with OP50 and supplemented with the following metabolites; Met – 5 mM methionine; FA – 100 µM folinic acid; Chol – 40 mM choline; 1 µM nucleosides. (H-K) Oocyte number of wild-type animals (H, J) and vulval induction index of *let-60(gf)* animals (I, K) fed with indicated RNAi clones or control (EV – empty vector); NGM was supplemented with 64 nM of B12; animals were exposed to RNAi from L1 except where indicated (L4). (B-K) Dots represent individual corpses/oocytes or animals (D, G, I, K) from two independent biological replicates; data represent median ± min and max; number of animals in brackets. For statistical analysis, see STAR methods. **** <0.0001; ***<0.0005; **<0.005; *<0.05; ns, non-significant.

To test if supplementing one-carbon metabolites reproduced the dietary effects on germ cells and VPC differentiation, we supplemented the growth medium with folinic acid, methionine and choline (bold in **Figure 1A)**. Wild-type animals fed with OP50 and exposed to one of the three metabolites had more apoptotic corpses and oocytes (**Figures 4B** and **4C**). Moreover, each of the three metabolites was sufficient to bypass the inhibitory effects of *metr-1(lf)* on germ cell differentiation (**Figures 4B** and **4C**). Only choline significantly enhanced VPC induction in *let-60(gf)* single mutants, but methionine and choline both reversed the effect of *metr-1(lf)* on vulval induction (**Figure 4D**). In contrast, folinic acid did not affect vulval induction (**Figure 4D**).

Since the folate cycle is involved in nucleotide production, we tested two bacterial strains that produce more nucleotides, *E. coli* OP50 cytR^-^zand *E. coli* HT115 (Chi et al., 2016). In the wild-type background, both strains enhanced germ cell apoptosis and oocyte differentiation (**Figures S5B** and **S5C**). Since *metr-1(lf)* worms were not viable when grown on these two bacterial strains, we examined somatically rescued *metr-1(lf)* mutants, which showed an increase in apoptotic corpse and oocyte numbers when fed with cytR^-^ or HT115 (**Figures S5B** and **S5C**), suggesting that nucleotide levels affect both processes. Vulval induction, on the other hand, was not enhanced by the two nucleotide-rich bacterial strains (**Figure S5D**). Furthermore, supplementing the growth medium with nucleosides caused an increase in apoptotic corpses and oocyte numbers in both wild-type animals and somatically rescued *metr-1(lf)* mutants (**Figures 4E** and **4F**), but did not affect vulval induction (**Figure 4G**). Similar to the DA1877 diet, supplementing choline or nucleosides partially suppressed the effects of germline-specific loss of *mpk-1* on germ cell apoptosis and oocyte development (**Figures S5E** and **S5F**), indicating that these metabolites enhance RAS/MAPK signaling. Taken together, folate and methionine cycle metabolites affect germ cell differentiation and death, whereas vulval induction is only sensitive to methionine cycle metabolites.

Phosphatidylcholine (PC) biosynthesis depends on choline (the Kennedy/CDP-choline pathway) and methyl groups provided by the methionine cycle to methylate phosphatidylethanolamine (PE) (**Figure 1A**). Since choline supplementation had a strong effect on all processes examined (**Figure 4B-D**), we hypothesize that PC levels might regulate germ cell and VPC differentiation. PC produced via the SAM-dependent pathway requires the *pmt-1* and *pmt-2* methyltransferases, while the choline-dependent pathway involves *cka-1* and *cept-1* activity (Brendza et al., 2007; Walker et al., 2011) (**Figure 1A**). We therefore used RNAi to test if the two PC synthesis pathways are necessary for the vitamin B12 effect on RAS/MAPK-controlled cell fates. Down-regulation of either *cka-1* or *cept-1* suppressed the vitamin B12-dependent increase in oocyte number and vulval induction (**Figures 4H** and **4I**). Due to the loss of the CED-1::GFP reporter signal in RNAi-treated animals, we could not score germ cell apoptosis. Since *pmt-2* RNAi induced larval arrest, we treated larvae with *pmt-2* RNAi bacteria diluted with empty vector (EV) starting at the L1 stage or exposed L4 larvae to undiluted *pmt-2* RNAi bacteria. In both treatments, *pmt-2* RNAi caused a reduction in oocyte number of wild-type animals supplemented with vitamin B12 (**Figure 4J**). *pmt-2* RNAi also reduced vulval induction in *let-60(gf)* animals supplemented with vitamin B12 (**Figure 4K**).

Together, these findings indicate that PC biosynthesis mediates the effects of vitamin B12 supplementation on oocyte and VPC differentiation, suggesting that the effects of choline are mediated by PC.

### The methionine cycle modulates VPC differentiation by repressing lipid metabolism

Since the one-carbon metabolism positively regulates germline and vulval development, and the DA1877 diet alters gene expression (Giese et al., 2020; MacNeil et al., 2013), we hypothesized that the DA1877 diet enhances germ cell and VPC differentiation by changing the expression of genes that modulate the cellular responses to RAS/MAPK signaling. RNAseq analysis of adult wild-type and *let-60(gf ^ts^)* animals fed with the OP50 or DA1877 diet identified in total 389 significantly down-regulated and 394 significantly up-regulated genes in DA1877-fed animals (**Figures 5A, 5B** and **Table S2**). In contrast to the metabolite levels, the DA1877 diet caused similar changes in the transcriptome of wild-type and *let-60(gf ^ts^)* worms (**Figure 5B**). Specifically, vitamin B12-, methionine cycle- and PC biosynthesis-related genes (*nhr-114, pmp-5, metr-1, sams-1, pmt-1* and *pmt-2*) were down-regulated in DA1877-fed worms (**Figures 5A, 5B** and **S6A**), suggesting a negative feedback response to increased levels of dietary vitamin B12 (Giese et al., 2020; MacNeil et al., 2013). Furthermore, we did not detect significant changes in the expression levels of RAS/MAPK pathway genes (**Figure S6A**). Pathway network analysis showed that the DA1877 diet regulates several genes involved in innate immune response and fatty acid biosynthesis (*fat-5*, *fat-6* and *fat-7*) (**Figure S6B**). Previous work has shown that diet-induced changes in gene expression can differ between young and gravid adult worms (MacNeil et al., 2013). We therefore tested if the DA1877 diet might influence gene expression in an age-dependent manner. We performed transcriptomic analysis of wild-type and *let-60(gf)* larvae at the mid-L3 stage, the stage when vulval induction occurs. The larval transcriptome showed smaller but similar changes in gene expression as observed in adults, as SAM, PC and fatty acid metabolism were also among the most strongly affected pathways in L3 larvae (**Figure S7A**-**C** and **Table S3**). We did not observe significant changes in *lin-1* and *lin-39* expression (**Figure S7C**), possibly because bulk RNAseq may not be sensitive enough to detect tissue-specific changes.

**Figure 5.**
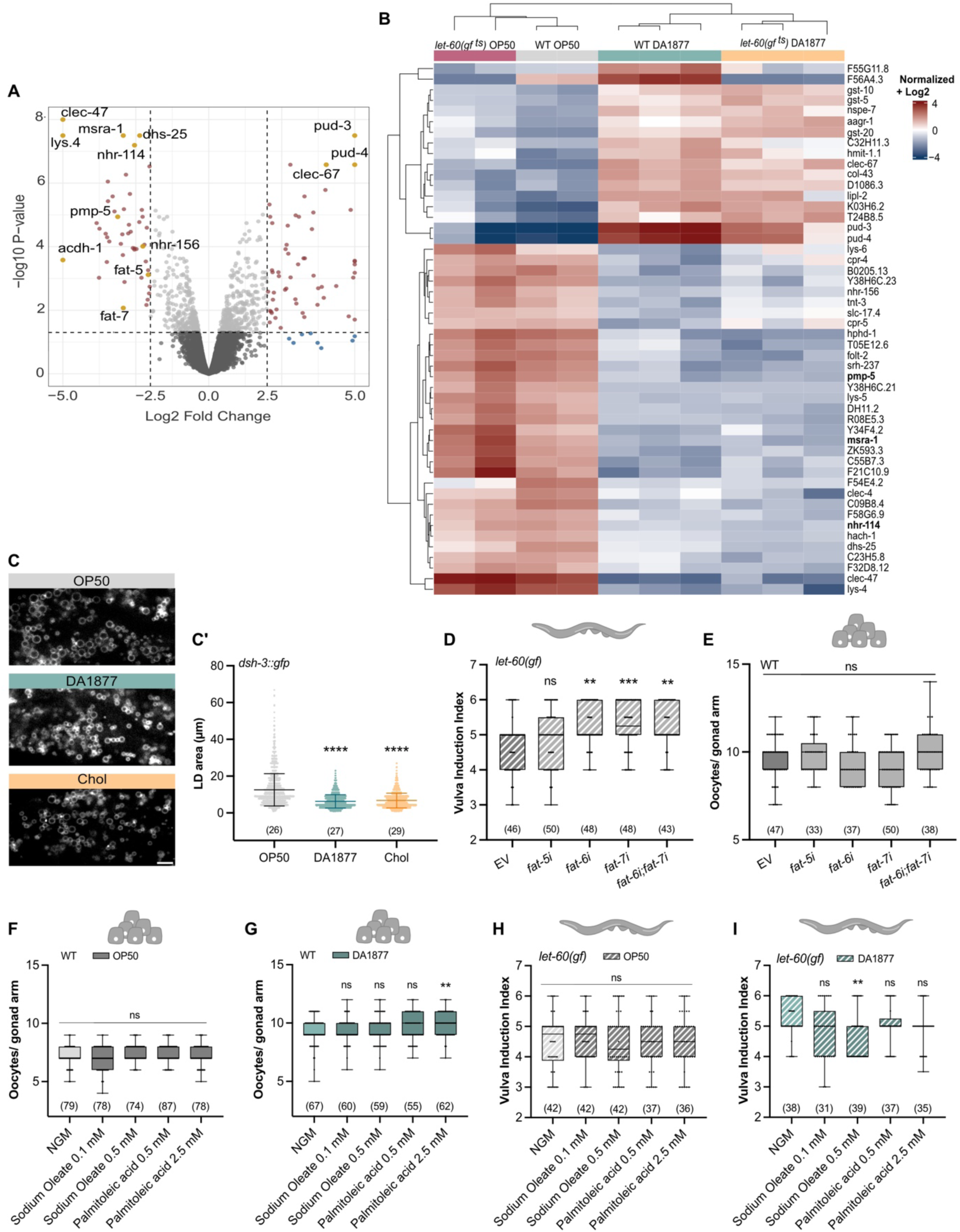
Dietary effects on gene expression and lipid metabolism. (A) Differentially expressed genes in adult wild-type animals fed with OP50 or DA1877; p-values calculated with FDR values and threshold for p-value 0.05 are shown. (B) Top 50 differentially expressed genes for indicated genotypes. (C) Fluorescence images of *dsh-3::gfp* marking intestinal lipid droplets (LD) in animals fed with OP50 (gray), DA1877 (blue) or OP50 supplemented with 40 mM choline (yellow). (C’) Area of intestinal LD; dots represent individual droplet areas of 100 droplets per animal, from two independent biological replicates. (D-E) vulval induction index of *let-60(gf)* animals (D) or oocyte number of wild-type animals (E) fed with indicated RNAi clones and control (EV – empty vector). (F-I) Oocyte number of wild-type animals (F, G) or vulval induction index of *let-60(gf)* animals (H, I) fed with OP50 (gray) (F, H) or DA1877 (blue) (G, I) supplemented with indicated fatty acids. (D-I) dots represent individual oocytes (E, F, G) or animals (D, H, I) from two independent biological replicates; data represent median ± min and max; number of animals in brackets. For statistical analysis, see STAR methods. **** <0.0001; ***<0.0005; **<0.005; ns, non-significant.

Previous work established a connection between vitamin B12, the methionine cycle and fat metabolism (Qin et al., 2022; Walker et al., 2011; Zhu et al., 2018). Given that genes controlling the lipid metabolism (i.e., the lipid Δ9-desaturases *fat-5*, *fat-6* and *fat-7*) were among the most strongly down-regulated genes, we explored the effect of the DA1877 diet on the lipid metabolism and its consequences on RAS/MAPK-induced cell fates. Quantification of lipid droplet size using the *dhs-3::gfp* reporter (Zhang et al., 2012) indicated that the intestines of DA1877-fed animals contained smaller lipid droplets than OP50-fed animals (**Figures 5C** and 5**C’**). Choline supplementation had a similar effect (**Figures 5C** and **5C’**), suggesting the DA1877 diet’s effect on lipid droplet size is choline-dependent. The *fat* genes are involved in different fatty acid desaturation and elongation pathways. *fat-5* is responsible for palmitoleic acid (C16:1n7) production, whereas *fat-6* and *fat-7* are responsible for oleic acid (C18:1n9) (Watts, 2009; Watts and Ristow, 2017). We tested if *fat-5, fat-6* or *fat-7* RNAi affected vulval induction and oocyte differentiation. Single or double knock-down of *fat-6* and *fat-7* increased vulval induction in *let-60(gf)* animals to a similar degree as the DA1877 diet, but *fat-5* knock-down had no significant effect (**Figure 5D**). Oocyte development was not affected by *fat-5, fat-6* or *fat-7* RNAi (**Figure 5E**). Finally, we supplemented palmitoleic acid and sodium oleate to animals fed with OP50 or DA1877. Oocyte differentiation was not chnged by fatty acid supplementation, irrespective of the diet (**Figures 5F** and **5G**). However, supplementing sodium oleate to *let-60(gf)* animals fed with the DA1877 diet reduced vulval induction to the levels observed in OP50-fed animals (**Figures 5H** and **5I**).

In summary, the down-regulation of unsaturated fatty acid biosynthesis by the DA1877 diet, especially oleic acid, positively regulated vulval induction, but it did not affect germ cell differentiation.

### A vitamin B12-rich diet increases global histone H3K4 tri-methylation to regulate VPC and oocyte differentiation

Since the methionine cycle is the main methyl donor in cells, we investigated whether changes in histone methylation might regulate germ cell and VPC differentiation. We focused on histone H3K4me3 because this type of histone modification is sensitive to diet, particularly a high-fat diet (Wan et al., 2022), as well as changes in methionine cycle activity (Ding et al., 2015, 2018; Godbole et al., 2023; Li et al., 2015). Western blot analysis showed higher global H3K4me3 levels in worms fed with DA1877 or supplemented with choline, and slightly reduced levels in *metr-1(lf)* mutants fed with OP50 (**Figure 6A**). Since the DA1877 diet down-regulated fat biosynthesis (**Figure 5C**), we tested if inhibiting *fat* genes affected histone methylation. H3K4me3 levels did not increase after *fat-6* or *fat-7* single or double RNAi, indicating that the oleic acid metabolism does not regulate H3K4me3 levels (**Figure 6B**).

**Figure 6.**
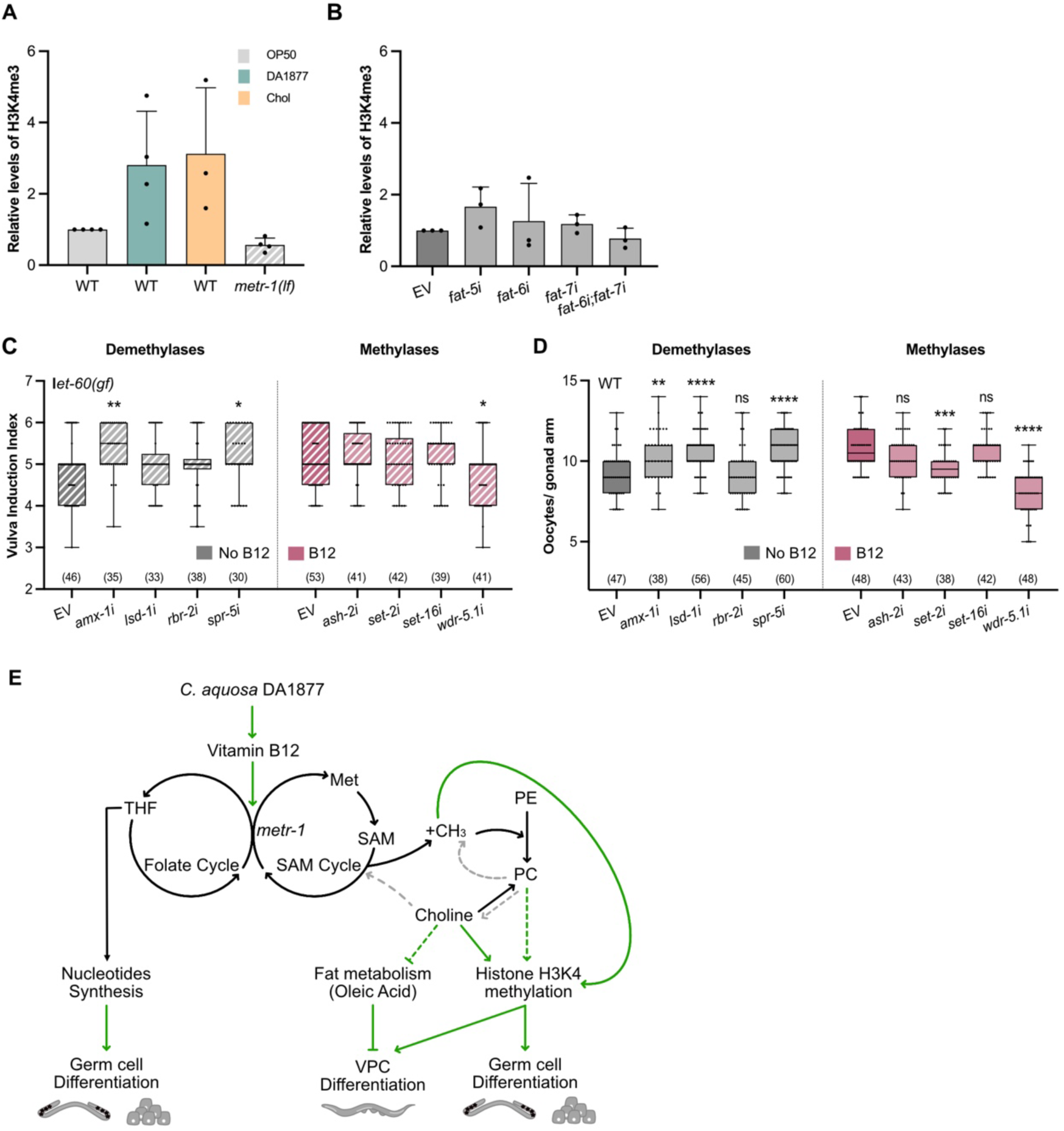
The DA1877 diet increases H3K4 methylation to regulate VPC and oocyte differentiation. (A) Western blot quantification of tri-methylated H3K4 for indicated genotypes fed with OP50 (gray), DA1877 (blue) or OP50 supplemented with 40 mM choline (yellow); levels of tri-methylated H3K4 were normalized to total H3 levels; bars represent mean ± SD, normalized to the control (wild-type OP50), from four independent biological and technical replicates. (B) Western blot quantification of tri-methylated H3K4 in wild-type animals fed with indicated RNAi clones; levels of tri-methylated H3K4 were normalized to total H3 levels; bars represent mean ± SD, normalized to the control (wild-type EV), from three independent biological and technical replicates. (C-D) Vulval induction of *let-60(gf)* animals (C) and oocyte number of wild-type animals (D) fed with indicated RNAi clones or control (EV – empty vector); non-supplemented NGM (gray) or supplemented with 64 nM B12 (pink); dots represent individual animals (C) or oocytes (D) from two independent biological replicates; data represent median ± min and max; number of animals in brackets. (E) Model depicting the tissue-specific effects of the one-carbon metabolism on nucleosides synthesis, fat metabolism and histone methylation and its effects on germ cell and VPCs differentiation; green arrows are interactions supported by this work; dashed arrows are hypothesized from literature (gray) or conclusions made in this work (green). For statistical analysis, see STAR methods. ***<0.0005; ***<0.0005; **<0.005; *<0.05; ns, non-significant.

To test if the increase in H3K4me3 levels caused by the DA1877 diet contributed to vulval induction and germ cell differentiation, we inhibited by RNAi the four H3K4 demethylases *amx-1, lsd-1, rbr-2* and *spr-5* (which might mimic the DA1877 diet in OP50-fed animals) and the four H3K4 methylases *ash-2, set-2, set-16* and *wdr-5.1* (which might suppress the vitamin B12 effect). RNAi of the demethylases *amx-1* and *spr-5* increased, and RNAi of the methylase *wdr-5.1* decreased vulval induction in *let-60(gf)* animals supplemented with vitamin B12 (**Figure 6C**). Oocyte differentiation was also increased by *amx-1, lsd-1* and *spr-5* RNAi and decreased by *set-2* and *wdr-5.1* RNAi (**Figure 6D**).

We conclude that a vitamin B12-rich diet causes a global increase in H3K4me3 levels, which contributes to the enhancement of RAS/MAPK-induced vulval induction and oocyte differentiation.

### Methionine-dependency of mammalian cells carrying activating KRAS and BRAF mutations

To investigate if the role of the methionine cycle in regulating RAS/MAPK-induced cell fates is conserved in mammals, we cultured human cancer cell lines carrying activating mutations in KRAS in medium containing varying methionine concentrations. We first performed collective cell migration assays with A549 lung cancer cells that carry a KRAS G12S mutation, to test if methionine restriction (MR) interfered with cell migration. To exclude possible effects of reduced cell proliferation, we performed these assays with cells arrested in the G1 phase by double thymidine blockade (Haag et al., 2020). A549 cells grown in MR (without methionine but supplemented with its precursor homocysteine) migrated at a lower speed than cells without MR (**Figures 7A, 7A’** and **7A’’**). The addition of the MEK kinase inhibitors MEK162 and LGX818 without MR reversibly blocked cell migration, indicating that RAS/MAPK signaling is essential for the migration of A549 cells (**Figure 7A, 7A’** and **7A’’**). Removal of the MEK inhibitors 8 hours after starting the assay allowed the cells to resume migration (**Figure 7A-A’**), indicating that the reduced migration after blocking the RAS/MAPK signaling was not due to increased cell death.

**Figure 7.**
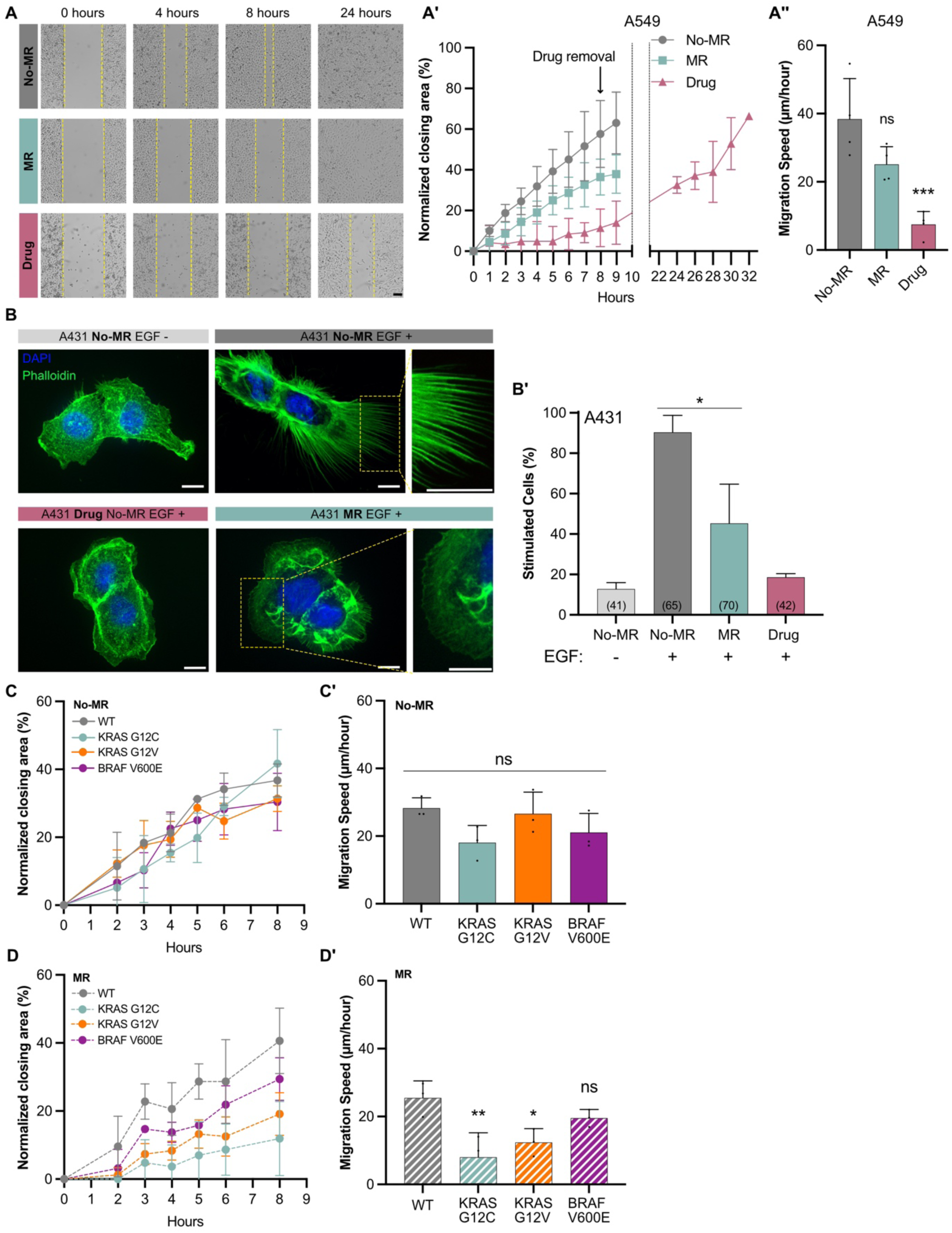
Methionine dependency of RAS/MAPK-induced phenotypes in mammalian cells. (A) Collective migration of A459 cells grown with MR (0 µM Met; 400 µM Hcy) (blue), without MR (200 µM Met; 400 µM Hcy) (gray), or treated with 1 µM MEK162 and 1 µM LGX818 MEK inhibitors (pink); yellow dashed lines represent the migrating front over time. (A’) Normalized percentage of the area closed over time; symbols indicate mean ± SD from four independent biological replicates. (A’’) Migration speed was calculated as the slope of graphs shown in A’; bars represent the mean ± SD. (B) Staining of A431 cells with DAPI (blue) and phalloidin (green), with and without EGF stimulation or MR; scale bar: 10 µm; for additional examples and the scoring criteria, see Figures S8. (B’) Percentage of stimulated cells with filopodia under indicated conditions; bars represent mean ± SD from three independent biological replicates; total number of cells scored in brackets. (C-D’) Quantification of collective migration in MEFs expressing different KRAS and BRAF mutations grown without (C) and with MR (D). (C-D) Normalized percentage of area closed over time; data represents mean ± SD from three independent biological replicates. (C’-D’) Migration speed was calculated as the slope of graphs shown in C and D respectively; bars represent the mean ± SD. For statistical analysis, see STAR methods. **<0.005; *<0.05; ns, non-significant.

We next used A431 epidermoid carcinoma cells, which overexpress EGFR and strongly respond to EGF stimulation by forming filopodia in a MAPK-dependent manner (Haag et al., 2020; Van de Vijver et al., 1991). Since filopodia formation is difficult to assess quantitatively, we categorized cellular morphologies into three classes; 0 for cells without filopodia, 1 for cells with small filopodia covering part of the cortex, and 2 for cells with long filopodia covering most of the cortex (**Figure S8A**). Class 1 and 2 cells were scored as stimulated and class 0 cells as unstimulated. Without EGF stimulation, most cells did not form filopodia, while 90% of the cells stimulated with EGF in the presence of methionine formed filopodia (**Figures 7B, 7B’** and **S8B**). By contrast, 55% of cells with MR and stimulated with EGF did not form filopodia (**Figures 7B, 7B’** and **S8B**), indicating that methionine is required for A431 cells to fully respond to EGF stimulation. Treatment with the MEK162 and LGX818 inhibitors in the presence of methionine strongly reduced EGF-induced filopodia formation (**Figures 7B and 7B’**).

Since cancer cell lines such as A459 carry many additional mutations besides the KRAS mutations, we used primary mouse embryonic fibroblasts (MEFs) in which the three RAS genes had been deleted and replaced with vectors expressing wild-type, G12C, G12V KRAS or V600E BRAF in the same genetic background (Drosten et al., 2010). Without MR, the KRAS mutant MEFs arrested in G1 showed the same migration speed as KRAS wild-type control cells (**Figures 7C and 7C**’). In contrast, KRAS G12V and G12C cells grown with MR migrated significantly slower, whereas KRAS wild-type MEFs migrated at the same speed with or without MR (**Figures 7D and 7D’**). V600E BRAF mutant MEFs showed a similar effect, though the reduction in migration caused by MR was smaller.

Taken together, these results suggested that the methionine-dependency of RAS/MAPK-induced phenotypes is conserved in mammalian cells.

## Discussion

Diet has a major impact on the cellular metabolism. Therefore, cells must adapt their metabolism to the available nutrition to ensure survival in changing environments. Here, we focused on the vitamin B12-dependent one-carbon metabolism and its effects on cell fate decisions controlled by the oncogenic RAS/MAPK signaling pathway in *C. elegans* (**Figure 6E**). A bacterial diet rich in vitamin B12 (DA1877) or direct vitamin B12 supplementation enhanced germ cell apoptosis, oocyte differentiation and VPC induction, which are all controlled by the RAS/MAPK pathway. Different one-carbon metabolites have tissue-specific effects, as the folate cycle only affected germline development, while methionine cycle metabolites enhanced all RAS/MAPK-induced phenotypes. Choline and phosphatidylcholine (PC) are key mediators of vitamin B12-dependent effects by regulating histone H3K4me3 methylation and repressing fatty acid biosynthesis (**Figure 6E**). We thus propose that methionine cycle activity is a limiting factor for different phenotypes caused by RAS/MAPK hyper-activation, by modulating the expression of downstream RAS/MAPK target genes. Finally, we show that methionine cycle activity is also a limiting factor in mammalian cells carrying activating mutations in KRAS and BRAF, pointing to a conserved role of the one-carbon metabolism in regulating cell RAS/MAPK-induced cell fates.

### The folate cycle as a tissue-specific regulator of germ cell differentiation

Folinic acid and nucleoside supplementation positively regulated germ cell death and oocyte differentiation, but not vulval induction. Both, the purine and pyrimidine metabolism are important for cell proliferation (Chi et al., 2016; Wan et al., 2019), which could explain our observations. For example, Chaudhari et al., 2016 showed that HT115 and DA1877 diets promote germline proliferation by increasing folate levels. Accordingly, DA1877- fed animals contained more mitotic germ cells in the proliferation zone and more M-phase cells. However, the DA1877 diet did not accelerate germ cell proliferation because the number of dividing cells relative to the total number of mitotic cells remained constant in a vitamin B12-rich diet. Moreover, the increase in oocyte numbers is not linked to germ cell apoptosis. However, the vitamin B12-rich diet accelerated germ cell transition through pachytene into diakinesis. While the mechanism by which nucleosides enhance germ cell apoptosis is not clear, we hypothesize that increasing nucleoside levels could promote oocyte differentiation by facilitating maternal mRNA production. Even though the most distal oocytes in diakinesis are transcriptionally silent, pachytene-stage germ cells are extremely active in producing mRNAs and proteins, which are taken up by the growing oocytes (Huelgas-Morales and Greenstein, 2018; Pazdernik and Sched, 2013; Walker et al., 2007). Since RAS/MAPK signaling promotes oocyte differentiation by regulating HTP-1 phosphorylation, it is also possible that a vitamin B12-rich diet regulates a meiotic checkpoint established by the HORMA complex (Das et al., 2020).

### The methionine cycle and phosphatidylcholine metabolism modulate RAS/MAPK signaling in all tissues

All RAS/MAPK-dependent cell fate decisions in *C. elegans* required an active methionine cycle, as evidenced by the suppression of the different *let-60 ras* phenotypes by *metr-1(lf)*. Interestingly, *let-60(gf)* animals fed with a vitamin B12-rich diet contained lower levels of methionine cycle metabolites than wild-type animals, while the levels of the transcripts encoding methionine cycle enzymes were not affected. It thus seems likely that *let-60(gf)* animals can produce these metabolites but consume them more rapidly, suggesting that hyper-activation of the RAS/MAPK pathway results in an increased dependency on methionine cycle metabolites.

The effects of the vitamin B12-rich diet are associated with increased choline and PC levels. PC biosynthesis and the RAS/MAPK pathway were previously connected in the context of uv1 cell survival under excitotoxic cell death (Crook et al., 2016). Hyper-activation of the RAS/MAPK pathway by LET-23 EGFR leads to uv1 cell survival, which requires PC biosynthesis. Additionally, choline, phosphocholine and PC levels rise in cancer cells because many enzymes of the choline metabolism are up-regulated by RAS/MAPK signaling (Glunde et al., 2011).

Choline supplementation not only increased vulval induction but also enhanced physiological germ cell death and oocyte differentiation. PC levels have previously been associated with apoptosis, with low levels of choline or PC-inducing apoptosis (Anthony et al., 1999; Cui and Houweling, 2002). However, apoptosis induced by choline deficiency is mediated by ceramide signaling and is mostly associated with stress- or radiation-induced apoptosis (Cui and Houweling, 2002; Yang et al., 2021; Yuan et al., 2021; Deng et al., 2008). In this context, our results point to a regulatory mechanism of physiological apoptosis controlled by choline metabolism. Given that PC is one of the most abundant phospholipids in cell membranes and that RAS/MAPK signaling is required for membrane integrity during oogenesis (Arur et al., 2011), choline could promote oocyte production by facilitating membrane production. This is consistent with our observation that germ cells in animals fed with the DA1877 diet exited pachytene faster, which may reflect a higher availability of membrane components.

### Fatty acid metabolism as a signaling pathway

A vitamin B12-rich diet induced the downregulation of *fat* genes. A relationship between the methionine cycle, choline and the lipid metabolism was previously reported (Li et al., 2011; Walker et al., 2011; Zhu et al., 2018). In particular, the methionine cycle and choline metabolism regulate lipogenesis in *C. elegans* via SBP-1, a homolog of the mammalian Sterol regulatory element-binding proteins (SREBPs) (Ding et al., 2015; Qin et al., 2022; Walker et al., 2011). RAS/MAPK signaling in cancer cells can also regulate SREBP activity (Glunde et al., 2011). These interactions are consistent with our data indicating that a vitamin B12-rich diet increases PC levels and reduces the size of lipid droplets in the intestine, probably by down-regulating *fat-5, fat-6* and *fat-7* expression. The relationship between PC and fat metabolism is not exclusive to *C. elegans*, as PC supplementation suppresses obesity-related phenotypes in mice fed with a high-fat diet (Lee et al., 2014), and a choline-deficient diet enhances liver fat levels in rats (Deminice et al., 2015).

Our data not only confirm the connection between the methionine cycle, choline and lipid metabolism but also show that fatty acid levels affect cell fate specification during vulval development. The fatty acid metabolism affects *C. elegans* in many ways, by regulating survival, developmental time, reproduction, ferroptosis and modulating drug effects (Brock et al., 2007; Diot et al., 2022; Goudeau et al., 2011; Lemieux and Ashrafi, 2016; Perez and Watts, 2021; Perez et al., 2020). However, our data point to specific functions of the different fatty acids, as only oleic acid inhibited VPC differentiation. Since fatty acids are known to act as signaling molecules (Dai et al., 2021; Watts and Ristow, 2017), we hypothesize that oleic acid regulates vulval development by activating a specific signaling pathway. Oleic acid is a monounsaturated fatty acid (MUFA) known for its antioxidant properties that stem from regulating pathways such as the p38 MAPK or the ROS pathway (Santa-María et al., 2023). In mice, oleic acid reduces kidney injuries by reducing inflammation via the RAS/MAPK/PPAR-y pathway (Zhang et al., 2022). Moreover, oleic acid has been reported to control apoptosis in rat hepatoma dRLh-84 cells (Yamasaki et al., 2008) and stimulate bovine neutrophils to function via intracellular calcium mobilization and ERK2 phosphorylation (Hidalgo et al., 2011).

### Methionine cycle, choline metabolism and histone methylation

A Vitamin B12-rich diet or choline supplementation increased global histone H3K4me3 levels, consistent with previous reports in worms and mammals indicating that the methionine cycle promotes H3K4 methylation (Dai et al., 2018; Kera et al., 2013; Mentch et al., 2015; Shyh-Chang et al., 2013). Also, choline can function as a methyl donor for DNA and histone methylation in mammals (Niculescu et al., 2006; Zeisel, 2011). Choline can be oxidized to betaine, providing a methyl group that is incorporated into homocysteine to regenerate SAM. However, the *C. elegans* genome does not encode an ortholog of choline oxidase (Wasmuth et al., 2008), and choline supplementation was not sufficient to bypass the reduction in H3K4me3 levels in intestinal cells caused by *sams-1* knock-down (Ding et al., 2015). Since phosphatidylethanolamine (PE) is a major consumer of methyl groups (Ye et al., 2017), SAM and thereby also H3K4me3 levels may increase if PC production is shifted to the Kennedy/CDP-choline pathway (**Figure 1A**). Thus, choline supplementation may promote histone methylation indirectly by reducing the competition for available methyl donors between histones and PE-dependent PC biosynthesis. Alternatively, choline could act directly as a methyl donor for H3K4 methylation.

H3K4me3 levels regulate different aspects of worm physiology such as fertility, lifespan and fat levels in *C. elegans* (Greer et al., 2010; Li and Kelly, 2011; Özdemir and Steiner, 2022; Xiao et al., 2011; Han et al., 2017). The H3K4 methyltransferases SET-16 and WDR-5.1 were previously reported to regulate VPC differentiation (Fisher et al., 2010). Here, we show that only WDR-5.1 is required to mediate the effects of dietary vitamin B12, while the demethylases the AMX-1 and SPR-5 appear to counteract WDR-5.1.

In summary, we propose that the methionine cycle is a limiting factor for RAS/MAPK-induced phenotypes in several tissues of *C. elegans*. Enhancing methionine cycle activity may modulate chromatin accessibility of specific target genes by increasing H3K4me3 methylation of target genes and thereby enhancing the cellular responses to RAS/MAPK signaling.

### Methionine-dependency of RAS/MAPK-regulated phenotypes in mammalian cells

The importance of RAS/MAPK signaling for oncogenic transformation is well documented, stimulating tumor cell proliferation, migration and invasion (Guo et al., 2020; Krueger et al., 2001; Schubbert et al., 2007). Also, the methionine-dependency of cancer cells has been known since 1959 (Sugimura et al., 1959). Non-carcinogenic cell lines can proliferate in the absence of methionine (MR), while many cancer cell lines show reduced or even blocked proliferation (Chello and Bertino, 1973; Sugimura et al., 1959). The exact reason why cancer cells need methionine is not completely understood. One possible explanation is that cancer cells have a reduced ability to use endogenous methionine for SAM synthesis due to a reallocation of homocysteine to the trans-sulfuration pathway (Cavuoto and Fenech, 2012; Kaiser, 2020). It is also possible that cancer cells have an increased demand for methionine and SAM due to more methylation reactions, rendering them more sensitive to MR (Cavuoto and Fenech, 2012; Booher et al., 2012; Kaiser, 2020). A methylation check-point where SAM levels regulate cell cycle progression through interactions with MCM, Cdc6 or Cdk2, has been proposed by Booher et al. (2012). Moreover, MR decreases global histone methylation, particularly H3K4me3 levels, which could down-regulate the expression of RAS/MAPK target genes (Kera et al., 2013; Mentch et al., 2015). The relationship between the one-carbon metabolism and cancer is used for cancer therapy, as multiple chemotherapeutic drugs such as 5-fluorouracil (5-FU) or methotrexate target the one-carbon metabolism (Ducker and Rabinowitz, 2017; Newman and Maddocks, 2017).

Here, we show that RAS/MAPK-regulated processes such as cell migration and filopodia formation depend on methionine. Using MEFs expressing different KRAS mutations as their only source of RAS activity, we observed remarkable differences in their sensitivity to MR. In the presence of methionine, KRAS and BRAF mutant MEFs were indistinguishable from wild-type MEFs but with MR, MEFs carrying activating KRAS mutations exhibited reduced cell migration. These findings support the idea of an increased dependency of RAS-transformed cancer cells on methionine cycle metabolites and mirror our findings with *C. elegans let-60 ras(gf)* mutants, which indicated that reducing methionine cycle activity limits the strength of different phenotypes caused by RAS/MAPK hyper-activation. MR in cancer cells may attenuate RAS/MAPK signaling through one or several of the mechanisms we discovered in the *C. elegans* model, i.e. H3K4me3 methylation, increased fat biosynthesis or reduced nucleoside production. Taken together, our findings suggest that the mechanisms regulating this key carcinogenic signaling pathway by the one-carbon metabolism may be conserved between worms and mammals.

## STAR Methods

### C. elegans culture and maintenance

*C. elegans* strains were maintained at 20°C on standard nematode growth medium (NGM) plates seeded with *Escherichia coli* OP50. *C. elegans* N2 (Bristol) strain was used as wild-type (Brenner, 1974). Before an experiment, worms were synchronized by bleaching gravid adults and letting the embryos hatch without food. *Comamonas aquosa* DA1877, *E. coli* OP50, *E. coli* HT115 and *E. coli* cytR^-^ were obtained from CGC. *C. aquosa* DA1877 ciA^-^/conB^-^ and *C. aquosa* DA1877 cbiB^-^ (Watson et al., 2014) were obtained from the Walhout lab. All bacterial strains were grown overnight at 37°C in liquid culture containing streptomycin (100 µg/ml) for DA1877, kanamycin (50 µg/ml) for cytR^-^ and streptomycin (100 µg/ml) and gentamycin (20 µg/ml) for both DA1877 mutants.

### Microscopy

Live *C. elegans* were mounted either on 3% agarose pads or in custom-made microfluidic devices for high-throughput and long-term imaging (Berger et al., 2018; Spiri et al., 2022). Nomarski (DIC) and epifluorescence images were acquired on one of three microscope systems: a DMRA2 (Leica) microscope equipped with a sCMOS camera (Prime BSI, Photometrics), a multicolor fluorescence light source (Spectra, Lumencor) and a piezo objective drive (MIPOS 100 SG, Piezosystems Jena); a DMRA (Leica) microscope equipped with two sCMOS cameras (C11440-42U30 Hamamatsu), an image splitter (TwinCam, Cairn Research), a multicolor fluorescence light source (Spectra, Lumencor) and a piezo objective drive (MIPOS 100 SG, Piezosystems Jena); or an inverted Ti-U (Nikon) equipped with a sCMOS camera (Prime 95B, Photometrics), a multicolor fluorescence light source (LedHub, Omicron Laserage Laserprodukt GmbH) and a piezo objective drive (Nano-F100, Mad City Labs). Images were acquired using a 20x air objective (HC PL APO 20x/0.80, Leica), a 40x oil objective (HCX PL APO 40x/1.32, Leica; CFI Plan Fluor 40X Oil, Nikon) or a 63x oil objective (HCX PL APO 63x/1.32-0.60, Leica). Confocal images were acquired on a spinning disk confocal (SDC) system consisting of a BX61 (Olympus) microscope equipped with an X-light V2 spinning disc unit, an EMCCD camera (iXon Ultra 888, Andor), two high-power LEDs (UHP-T-460-DI and UHP-T-560-DI, Prizmatix) for GFP and RFP excitation, a mercury vapor lamp (X-Cite exacter, Excelitas Technologies Corp) for DAPI, CFP and YFP excitation and a piezo objective drive (MIPOS 100 SG, Piezosystems Jena). Images were acquired using a 40x oil objective (UplanFL N 40x/1.30, Olympus) or a 60x oil objective (UPlanAPO 60x/1.40, Olympus).

### Germ cell apoptosis and oocyte numbers

Unless stated otherwise, all experiments were performed in one-day-old adult animals (72 hours post L1 starvation) grown at 20°C. To score germ cell apoptosis in temperature-sensitive *let-60(ga89) (gf ^ts^)* animals, all worms were initially grown at 15°C and transferred to 25°C for 4 hours before corpse scoring. Germ cell apoptosis was scored using the *bcIs39[Plim-7::ced-1::gfp]* reporter (Derry et al., 2001) and quantified in one gonad arm by counting CED-1::GFP positive cells. To count oocytes with the *let-60(ga89) (gf ^ts^)* allele, worms were initially grown at 20°C and transferred to 25°C for 18 hours before counting oocyte numbers. Oocytes were scored in DIC images or using the SYN-4::GFP membrane marker, counting the number of completely cellularized oocytes in the proximal gonad arm.

### Vulval induction

All vulval induction experiments were conducted in the *let-60(n1046^gf^)* background. Vulval induction was scored in L4 animals (46-48 hours post-L1) using DIC images (Sternberg and Horvitz, 1986). The vulval induction index, the average number of induced VPCs per animal, was calculated by assigning a score of 1 to a VPC that had undergone three rounds of cell divisions, and a score of 0.5 when only one of the two VPC descendants had differentiated.

### Metabolites, nucleoside and fatty acid supplementation

Vitamin B12, metabolites and nucleosides were added to freshly prepared NGM plates. Vitamin B12 was dissolved in water at 1 mM stock concentration and added at a final concentration of 64 nM unless indicated otherwise. All of the metabolites were directly added to the NGM plates at the following final concentrations: 40 mM for choline, 5 mM for L-methionine, and 100 µM for folinic acid. Nucleosides were also added directly at a final concentration of 1 mM each. Sodium oleate was likewise added directly at a final concentration of 0.1 and 0.5 mM, and palmitoleic acid at a final concentration of 0.5 and 2.5 mM. For fatty acid supplementation, 1 µM tergitol (NP40S) was included in the final mix for solubilization (Deline et al., 2013). For suppliers and catalog numbers, see **Key Resource Table**.

### Long-term image acquisition

Long-term imaging experiments were performed as described by Berger et al., 2018. Briefly, OP50 or DA1877 bacteria were grown overnight at 37°C in LB medium. Bacterial cultures were then washed three times with S-Basal buffer and concentrated to 1 ml. For OP50, the bacterial culture was re-suspended in 650 µl of Optiprep and 332 µl of S-Basal supplemented with 1% Pluronic F127 and for DA1877 in 600 µl of Optiprep and 380 µl of S-Basal supplemented with 1% Pluronic F127. The bacterial suspension was filtered using a 10 µm strainer (pluriStrainer Mini 10 um, PluriSelect). Worms were washed three times with S-Basal before loading on the chip. Animals were loaded as described, and immobilized by the microfluidic device channel and two on-chip hydraulic valves (Berger et al., 2018). Epifluorescence images were acquired using a 40x oil objective, at an interval of 10 minutes for germ cell tracking, and 5 minutes to measure ovulation rates. Wild-type animals were imaged at 20°C, and *let-60(ga89)* animals at 25°C, with animals shifted from 20°C to 25°C two hours before the start of the experiment. The temperature was controlled (±0.5°C) by either the room’s air conditioning system or a microscope cage incubator (H201-T-UNIT-BL-CRYO and H201-ENCLOSURE-CRYO, Okolab).

### Germ cell tracking

Worm movement during long-term imaging acquisition was corrected using a custom-built MATLAB script. Briefly, sample drift in the z-direction was compensated for each time point assuming linear sample drift within the device channel. Image registration was initially performed on the channel outlining the germ cell membrane, with the derived transformation applied to all other channels. Sample shift in between time points was compensated by visually identifying the gonad’s loop region and cropping the original image to a final size of 750x250 pixels from the loop. For each time point, a maximum intensity z-projection of 5-10 slices was created.

Germ cell migration was tracked over 2 hours, manually identifying the position of individual germ cells in each projection. Germ cells were tracked using the SYN-4::GFP membrane marker alone for *let-60(ga89)(gf ^ts^)* or combined with the mCherry::H2B marker for the wild-type. Since germ cells migrate collectively, at least 10 individual cells distributed over the distal gonad arm were tracked per worm, and eight worms per genotype were analyzed.

### RNAi Interference

RNAi plates were prepared by adding ampicillin (100 µg/ml) and IPTG (1 mM) to the NGM mixture. *E. coli* HT115 RNAi cultures were grown overnight at 37°C with shaking in LB medium supplemented with ampicillin (100 µg/ml) and tetracycline (10 µg/ml). Before plating the bacteria on the RNAi plates, the overnight liquid culture was diluted 1:1 with fresh LB (supplemented with ampicillin, tetracycline and IPTG) and bacteria were allowed to grow for 3-5 hours at 37°C with shaking (except for the *pmt-2* RNAi clone). 400 µl of bacterial culture were seeded on each plate, and plates were incubated at 37°C overnight. RNAi clones were obtained from either the Ahringer RNAi library or the ORFeome RNAi library (both from Source BioScience). All RNAi clones were sequenced before the experiments.

### ERK-nKTR biosensor quantification

MPK-1 activity in the VPCs was measured in mid-L2 larvae (1-cell-stage) using the ERK-nKTR biosensor described in de la Cova et al., 2017. Worms fed on OP50 were imaged 20-23 hours post L1 and worms fed on DA1877 were imaged 19-21 hours post L1. The developmental stage was determined by measuring gonad length (35-75 µm in mid-L2). Epifluorescence images were acquired with a 0.25 µm z-step, using a 63x oil objective described as above.

Image analysis was performed using a custom-built MATLAB script. Acquired images were flat-field corrected, after which the in-focus slice was identified using the mCherry::H2B nuclear signal. The images were then z-projected (sum intensity) around the in-focus slice (two slices above and below) and individual VPCs were identified. The red/green fluorescence intensity ratio was calculated for each VPC except P3.p, and normalized to the average intensity ratio of all VPCs for each animal.

### Mitotic Index of germ cells

The mitotic zone of dissected gonads was identified by HIM-3 antibody staining (see Gonad dissection and immunostaining for sample preparation). Mitotic cells were counted using Fiji software (Schindelin et al., 2012) based on nuclear DAPI staining. The number of pH3-positive M phase nuclei was divided by the total number of mitotic cells to calculate the mitotic index. Images were acquired using a confocal microscope with a 60x objective.

### Sample preparation for LC-MS analysis

Approximately 3000 worms were washed three times with ice-cold M9. M9 was then completely removed to obtain a pellet of adult worms, which was snap-frozen in liquid nitrogen and stored at −80°C until all three biological replicates had been collected. Sample preparation and mass spectrometry analysis were performed by the Functional Genomics Center Zurich (FGCZ). For tissue homogenization, *C. elegans* larvae were placed in test tubes with 1 ml precooled MCC (MeOH:CAN:Water 4:4:2) and one precooled 5 mm stainless steel bead. Homogenization and metabolite extraction were achieved by performing two rounds of 2.5 min of shaking at 30 Hz (with precooled tubes and racks) using a TissueLyser (Qiagen). Homogenates were centrifuged for 10 min at 10’000 rpm/4°C and metabolite extracts were recovered to clean test tubes. Protein concentration was measured in the extraction pellet and used for data normalization.

Before measurement, extracts were reconstituted in an injection buffer (90% acetonitrile). The solution was vortexed and centrifuged for 10 min at 12’000 rpm/4°C and 60 μl of the supernatant was transferred to a glass vial with a narrowed bottom (Total Recovery Vials, Waters) for LC-MS injection. In addition, method blanks, QC standards, and pooled samples were prepared in the same way to serve as quality control for the measurements.

### LC-MS analysis

Metabolites were separated on a Thermo Vanquish Horizon Binary Pump equipped with a Waters Premier BEH Amide column (150mm x 2.1mm) by applying a gradient of 10 mM ammonium bicarbonate in water (A) and 10 mM ammonium bicarbonate in 95% acetonitrile (B) from 99% B to 30% B over 12 min. The injection volume was 3.5 μl. The flow rate was 0.4 μl/min with a column temperature of 40°C and autosampler temperature of 5°C. The LC was coupled to a Thermo Qexactive mass spectrometer by a HESI source. MS1 (molecular ion) and MS2 (fragment) data were acquired using negative polarization and Full MS / dd-MS² (Top5) over a mass range of 70 to 1050 m/z at MS1 and MS2 resolution of >17’500.

### Untargeted Metabolomics Data analysis

Metabolomics data sets were evaluated in an untargeted fashion with the Compound Discoverer software (Thermo Scientific), which aligns the ion intensity maps and performs peak picking on an aggregated ion intensity map. Detected ions were identified based on accurate mass, MS2 data, detected adduct patterns and isotope patterns by comparing with entries in the online databases and in-house analyzed collection of standards (IROA MSMLS kit, IROA Technologies™). Quality controls were run on pooled samples and reference compound mixtures to determine technical accuracy and stability.

### mRNA isolation and sequencing

For transcriptome analysis by RNAseq, animals were collected from plates seeded with either OP50 or DA1877 bacteria. For adults, approximately 500 animals (wild-type or *let-60(ga89) (gf ^ts^)*) were collected. For larval analysis, 1500 animals (wild-type or *let-60(n1046^gf^)*) were collected. Before sample collection, adult worms were transferred to 25°C for 4 hours. Three biological replicates were used per condition. Animals were washed off plates with M9 and snap-frozen in liquid nitrogen. Samples were stored at −80°C until RNA extraction. Total mRNA extraction was performed for all samples at the same time with trizol, and samples were purified using a Rneasy Mini Kit (QIAGEN). Quality control (Agilent DNF-471 RNA kit), cDNA library preparation and whole exome sequencing (Illumina Novaseq 6000 system) were performed by the transcriptomics service of the Functional Genomics Center Zurich (FGCZ). Between 15 and 30 million reads were obtained per sample. Bioinformatic analysis was performed using an FGCZ in-house bioinformatics pipeline using the open-source tools STAR for read alignment and DESeq2 for differential gene expression.

### Western blot analysis

One hundred worms were washed three times in ice-cold M9 and lysed in 50 µl 2x SDS PAGE buffer (containing 200 mM β-MCE) for 5 minutes at 95°C. Genomic DNA was digested by adding 2 µl of Rnase-free Dnase (QIAGEN) for 5 minutes at room temperature followed by a 5-minute incubation at 95°C. Samples were kept at 4°C overnight or −20°C for longer periods before loading. Before loading, samples were heated to 95°C, and 15 µl were loaded onto 4-12% gradient polyacrylamide gels.

For MAPK, histone H3 and histone H3 tri methyl K4 quantification, the membrane was treated with transfer buffer containing 20% methanol and blocked with 5% BSA for anti-MAPK and 5% milk for anti-histone blots. Samples were incubated with primary antibodies at 4°C overnight. Secondary antibodies (1:2000) HRP conjugated anti-rabbit or HRP conjugated anti-mouse were incubated for 2 hours at RT. The HRP was detected by incubating the membrane with a chemiluminescence assay for 4 minutes. For MAPK quantification, the membrane was then incubated with sodium azide (1:10) for 1 hour to inactivate the HRP-conjugates, and re-stained for alpha Tubulin (1:10’000), followed by incubation with HRP-conjugated secondary antibodies, as well as the chemiluminescence assay. The following primary antibodies were used: 1:2000 anti-dephosphorylated ERK 1&2 Mouse; 1:10’000 anti-MAP Kinase Rabbit; 1:1000 anti-histone H3 rabbit and anti-histone H3 tri methyl K4. Signal levels were quantified by measuring band intensities using a built-in tool in Fiji (Schindelin et al., 2012). For MAPK quantification, both total MAPK and phospho-MAPK were normalized to the tubulin loading control. Phospho-MAPK levels were then normalized by total MAPK levels.

### Gonad dissection and immunostaining

Gonad dissection was performed in PBS containing 0.2 mM tetramisole. Dissected gonads were transferred to siliconized 2 ml Eppendorf tubes and fixed in 4% formaldehyde for 10-60 minutes. For HIM-3 and pH3 staining, gonads were washed three times with PBS-T (0.05% Tween 20) and permeabilized with 0.5% Triton for 12 minutes. All gonads were washed three times with 0.05% PBS-T and post-fixed for 5 minutes at −20°C in 100% methanol. Dissected gonads were blocked for 2-4 hours at room temperature in PGB buffer (PBS-T, 1% BSA, 0.2% gelatin) or 1% BSA. Primary antibodies were diluted in PGB buffer and incubated overnight at 4°C, except for pH3 staining, which was incubated for 4 hours at room temperature after primary incubation with HIM-3 overnight at 4°C. Secondary antibodies were incubated for 2 hours at room temperature. After secondary incubation, gonads were washed three times with PBS-T. To stain germ cell nuclei DAPI dihydrochloride was added (1:10’000) during the second wash for 10 minutes. Gonads were mounted in Mowiol on a glass slide and polymerized overnight at 4°C. Epifluorescence images were acquired using a 20x air objective as described above. Before quantification, images were flat-field corrected using a custom-built MATLAB script. To correct for differences in gonad length, the length of the distal arm was normalized such that the distal tip cell (DTC) corresponded to 0 and the loop region to 1. Intensity profiles from the DTC to the loop were measured along each gonad in sum-intensity projections using Fiji (Schindelin et al., 2012).

**Table 1:** Antibodies used for immunostaining.

| Antibody | Dilution |
| --- | --- |
| Anti-dephosphorylated ERK 1&2 Mouse | 1:2000 |
| Anti-XND-1 Guinea-Pig | 1:2000 |
| Anti-SUN-1 (pS8) Guinea-Pig | 1:700 |
| Anti-GLD-1 Rabbit | 1:100 |
| Anti-HIM-3 Rabbit | 1:500 |
| Anti-phospho-histone H3 (Ser10) | 1:400 |
| Anti-Rabbit Alexa 568 Donkey | 1:1000 |
| Anti-Guinea Pig Alexa 594 Goat | 1:1000 |
| Anti-mouse Alexa 488 Goat | 1:1000 |

### Quantification of reporter gene expression

LIN-1::GFP and LIN-39::GFP fluorescence intensity were scored in VPC descendants at the 2-cell Pnp.x stage (mid-L3). Wild-type worms fed with OP50 were imaged 29-30 hours post L1; wild-type worms fed with DA1877 were imaged 27-29 hours post L1; *let-60(n1046) (gf)* worms fed with OP50 were imaged 32-33 hours post-L1; and *let-60(n1046) (gf)* worms fed with DA1877 were imaged 29 hours post L1. Images were acquired on an epifluorescence microscope using a 63x oil-immersion lens, as described above. Before quantification, images were flat-field corrected using a custom-built MATLAB script. VPC nuclei were manually selected and the average fluorescence intensity was measured on an in-focus z-slice using built-in tools in Fiji (Schindelin et al., 2012). Fluorescence intensities for each pair of VPC descendants were averaged. The data for each VPC was normalized to the average value obtained from OP50-fed animals (i.e., P5.p on DA1877 or OP50 divided by the average of P5.p on OP50).

### Quantification of lipid droplet size

Lipid droplet size was measured using the DHS-3::GFP reporter (Zhang et al., 2012). Twenty worms were analyzed per condition, and the area of 100 droplets was measured in the posterior section of the intestine of each worm. Images were acquired using a confocal microscope using a 100x oil-immersion lens as described above. Lipid droplet area was measured by manually selecting lipid droplets using Fiji (Schindelin et al., 2012).

### Generation of transgenic C. elegans

Somatic rescue of *metr-1* was achieved by creating extrachromosomal arrays in the *metr-1(lf)* background. Microinjections were performed according to Mello et al., 1991. Purified PCR DNA was injected at a concentration of 20 ng/µl. Two co-injection markers were used: 2.5 ng/µl of pCFJ90 (*Pmyo-2>mCherry*) (Frøkjaer-Jensen et al., 2008) and 100 ng/µl of pBluescript-KS. Three independent lines were generated and characterized.

### Mammalian cell culture

For cell culture experiments, we used the human lung carcinoma epithelial cell line A459, the epidermoid carcinoma A431 cell line and four mouse embryonic fibroblast (MEFs) lines obtained from the National Cancer Institute (NIH): KRAS 4B WT, KRAS 4B G12C, KRAS 4B G12V and BRAF V600E. All cell lines were tested for mycoplasma and kept at 5% CO_2_, 98% relative humidity and 37°C in a modified DMEM medium with high glucose and sodium pyruvate, containing 10% fetal bovine serum (FBS).

For methionine restriction (MR) experiments, cells were kept in a modified DMEM medium with high glucose. For methionine-positive conditions (no-MR), the medium was supplemented with 200 mM L-Glutamine, 200 µM L-Methionine, 100 mM L-Cystine dihydrochloride, 400 µM DL-Homocysteine thiolactone hydrochloride and 100 mM sodium pyruvate. For methionine-restricted (MR) conditions, the medium was equally supplemented except for methionine.

### Cell migration assays

G1-arrested A549 cells and MEFs were used for migration assays. Cells were seeded at 80’000 cells/ml density in DMEM medium with or without supplements. After 32 hours, cells were exposed to thymidine (1:50) to arrest the cell cycle as described in Haag et al., 2020 (first thymidine block). 16 hours after, the first thymidine block was released by replacing the medium. 8 hours after, cells were transferred to ibidi 2-well silicon culture inserts at a density of 400’000 cells/mL and again exposed to a medium containing thymidine (second thymidine block). Cells remained in the silicon culture inserts for 16 hours after which the inserts were removed and pictures were taken every hour for 8 hours. For experiments with MEK inhibitors, cells were exposed to 1 µM MEK162 (binimetinib) and 1 µM LGX818 (ecorafenib) during the first thymidine block. The relative migration was calculated as *R[%]=(R0—Rn)/R0×100[%],* where R0 represents the initial open area and Rn represents the remaining open area at each time point. Linear regression was used to determine migration rates shown in the bar graphs (**Figure 7A’’**).

### EGF stimulation and immunostaining of A431 cells

The A431 cell line was used to analyze filopodia formation upon EGF stimulation. Cells were exposed to no-methionine restriction (no-MR, 200 µM methionine + 400 µM homocysteine) or MR (0 µM methionine + 400 µM homocysteine) 24 hours before the experiment. Cells were then seeded on glass slides in 24-well plates at 114’000 cells/ml density in no-MR or MR conditions. After 24 hours, cells were serum starved and treated with 1 µM MEK162 (binimetinib) and 1 µM LGX818 (ecorafenib). After another 16 hours, cells were stimulated with 100 ng/ml of EGF for 10 minutes at 37°C (Haag et al., 2020), followed by 15 min fixation in 4% PFA at 37°C. Cells were permeabilized for 5 min with 0.2% Triton X-100% and 0.5% BSA and then blocked for 1 hour in 0.5% BSA and 0.2% gelatine. Fixed cells were stained with Phalloidin 568 for 40 minutes and 0.1 µg/ml DAPI for 5 minutes. Glass slides were mounted with ProLong Gold Antifade Mountant (Thermo Scientific).

### Statistical analysis

Statistical analysis was performed using GraphPad Prism 10. When comparing two samples, an unpaired t-test was used. For multiple samples, data was analyzed by one-way ANOVA followed by Dunnett’s multiple comparison test for parametric data and a Kruskal-Wallis test for non-parametric data.

## Acknowledgments

We would like to thank all present and past members of the Hajnal group for critical discussion and input on this project. A special thanks to Michael Daube for lab assistance and maintenance and to Dr. Michael Walser for support during the cell culture. We are thankful to the *Caenorhabditis* Genetics Center (CGC) (funded by the NIH P40 OD010440), to Wormbase (funded by NIH RO1 OD023041), Dr. Walhout and her lab for sending us the *C. aquosa* vitamin B12-mutants, and the National Cancer Institute (NIH) for the KRAS mutant and wild-type MEFs. We also want to thank Drs. Verena Jantsch, Judith Kimble and Monique Zetka for providing antibodies. We thank Dr. deMello and his lab for access to their facilities and reagents. Finally, a big thank you to the Functional Genomics Center Zurich (FGCZ) of the University of Zurich, especially Dr. Alaa Othman and Dr. Martina Zanella for helping us with the metabolomic analysis and Dr. Maria Domenica Moccia for help with the RNAseq. This project was supported by the Candoc Forschungskredit of the University of Zurich to A.C.L. (grant no. K-74406-03) and the Swiss National Science Foundation (grant no.31003A-184792) to A.H.

## Author contribution

Conceptualization: A.C.L. and A.H.; Investigation: A.C.L., S.B., T.K. and A.H.; Formal Analysis: A.C.L., S.B. and N.R.G.; Writing – Original draft: A.C.L. and A.H.; Writing – Review and Editing: A.C.L. and A.H.; Funding acquisition: A.C.L. and A.H.

## Supplementary Figures and Legends

**Figure S1.**
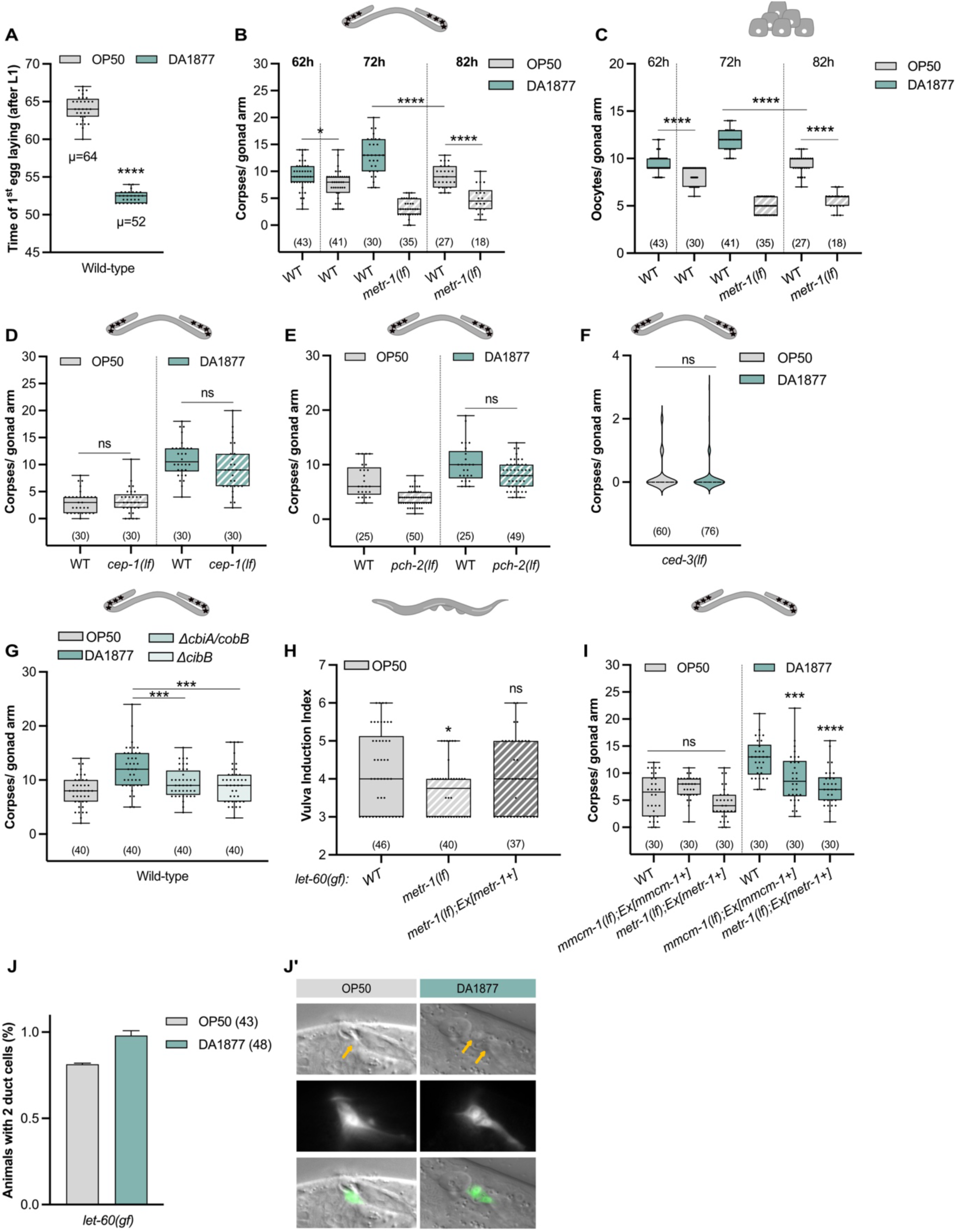
Dietary vitamin B12 effect on germline, vulva and duct cell specification. (A) Time of first egg laying after L1 in wild-type worms fed with OP50 (gray) or DA1877 (blue). (B-C) Number of corpses (B) or oocytes (C) for indicated genotypes 62, 72 and 82 hours after L1 starvation; animals were fed with OP50 (gray) or DA1877 (blue). (D-F) Number of corpses for indicated genotypes fed with OP50 (gray) or DA1877 (blue). (G) Number of corpses in wild-type animals fed with OP50 (gray), wild-type DA1877 (blue) or DA1877 mutants deficient in B12 synthesis (light blue). (H) Vulval induction index for indicated genotypes fed with OP50. (I) Number of corpses for indicated genotypes fed with OP50 (gray) or DA1877 (blue). (A-I) Data represent median ± min and max. (J) Percentage of *let-60(gf)* animals with two duct cells, animals were fed with OP50 (gray) or DA1877 (blue); bars represent mean ± SD. (J’) DIC and fluorescence images of *let-60(gf)* animals expressing the LIN-48::GFP duct cell marker.; yellow arrows indicate the duct cell; scale bar: 10µm. (A-G, I) dots represent individual corpses/oocytes; (H) dots represent individual animals; number of animals in brackets. For statistical analysis, see STAR methods. **** <0.0001; ***<0.0005; *<0.05; ns -non-significant.

**Figure S2.**
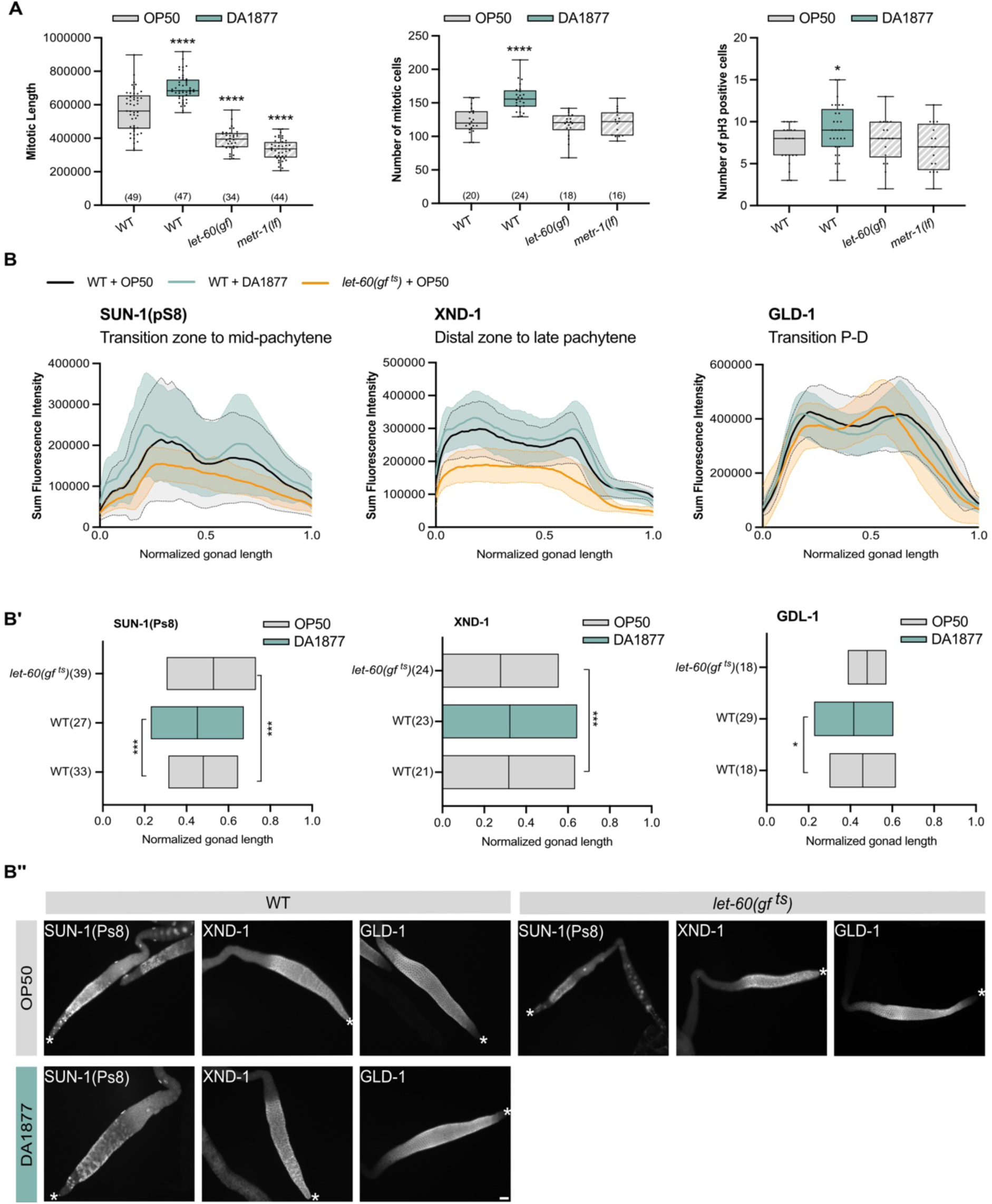
Effects of the DA1877 diet on germline dynamics. (A) Length of the mitotic zone, number of mitotic cells and number of M-phase germ cells (pH3 positive) for indicated genotypes fed with OP50 (gray) or DA188 (blue); dots present individual animals; data represents median ± min and max; number of animals represented in brackets. (B) Summed intensity profile of SUN-1(pS8), XND-1 and GLD-1 staining along the distal-proximal axis of dissected gonads. In wild-type animals fed with OP50 (black), SUN-1(pS8) accumulates from the transition zone until mid-pachytene; XND-1 starts at the distal tip and accumulates until late-pachytene; GLD-1 marks the transition between pachytene and diplotene. Animals were fed OP50 (gray) or DA1877 (blue), and *let-60(gf ^ts^)* animals were fed on OP50 (orange) and transferred to 25°C 2 hours before the experiment. The x-axis represents the normalized distance along the gonad, from distal (x=0) to the loop (x=1); lines indicate average intensity ± SD (shaded). (B’) Analysis for SUN-1, XND-1 and GLD-1 immunostainings. Graphs represent the location of the maximum intensity peaks for the distal (beginning of expressions) or proximal (end of expression) part of the gonad (from B); the x-axis represents the normalized distance; bars represent median ± min and max. (B’’) Examples of immunostaining of dissected gonads of wild-type animals fed with OP50 (gray) or DA1877 (blue), and *let-60(gf ^ts^)* animals fed with OP50 (gray); white stars indicate the distal tip cell; scale bar: 10 µm. For statistical analysis see STAR methods. **** <0.0001; ***<0.0005; *<0.05.

**Figure S3.**
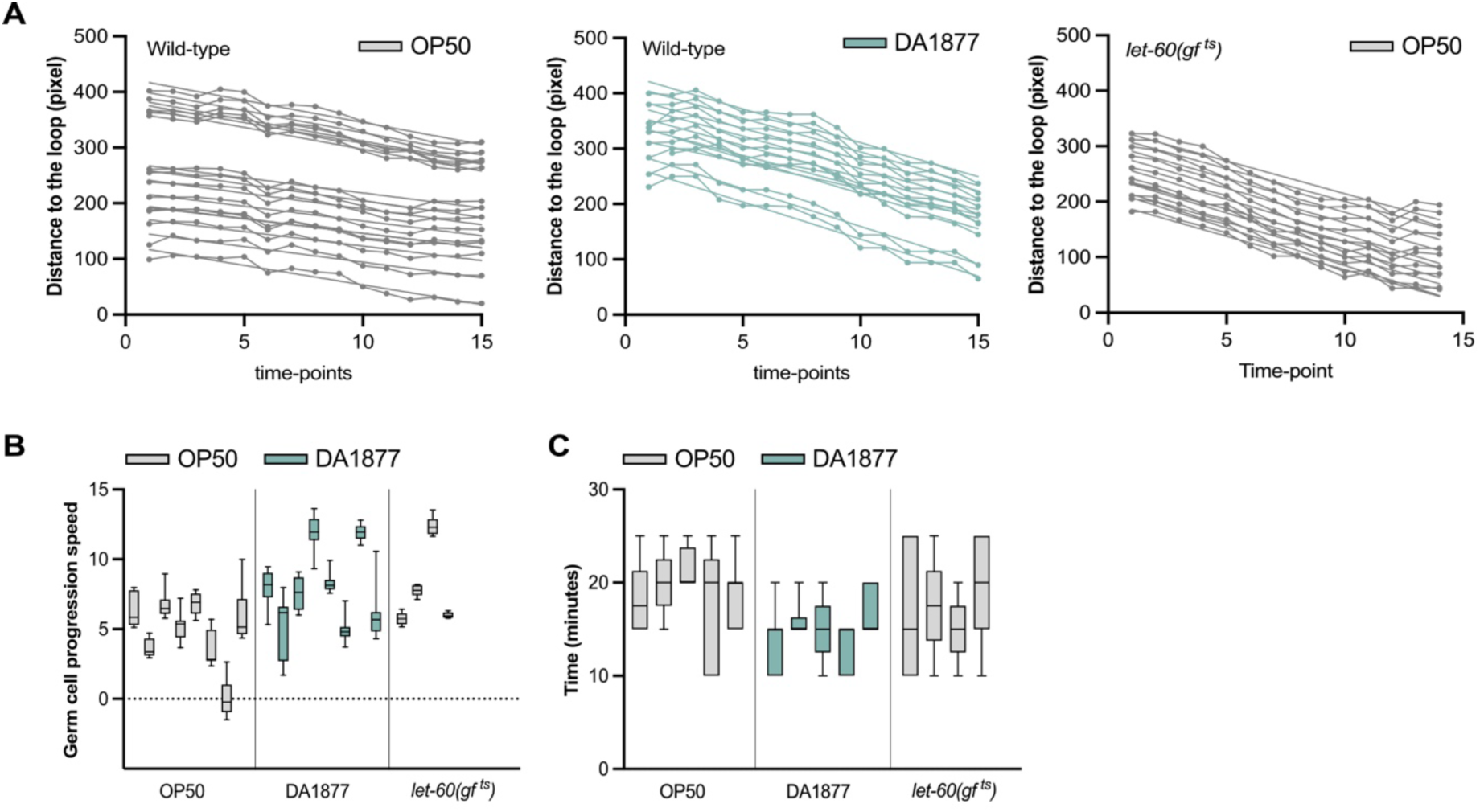
Effects of the DA1877 diet on germ cell migration and ovulation rate. (A) Tracking of individual germ cells within the pachytene region over time for indicated genotypes; pachytene stage germ cells before the loop were tracked for 2 hours; migration speed was approximated for each graph as the slope of a linear regression, shown in Figure 2G. The graphs represent cells from one of eight animals tracked for the wild-type and four for *let-60(gf ^ts^)* animals that were transferred to 25°C 2 hours before the experiment and imaged at 25°C. (B) Speed of tracked germ cells for individual animals. (C) Ovulation time for individual animals. (A-C) animals were fed with OP50 (gray) or DA1877 (blue).

**Figure S4.**
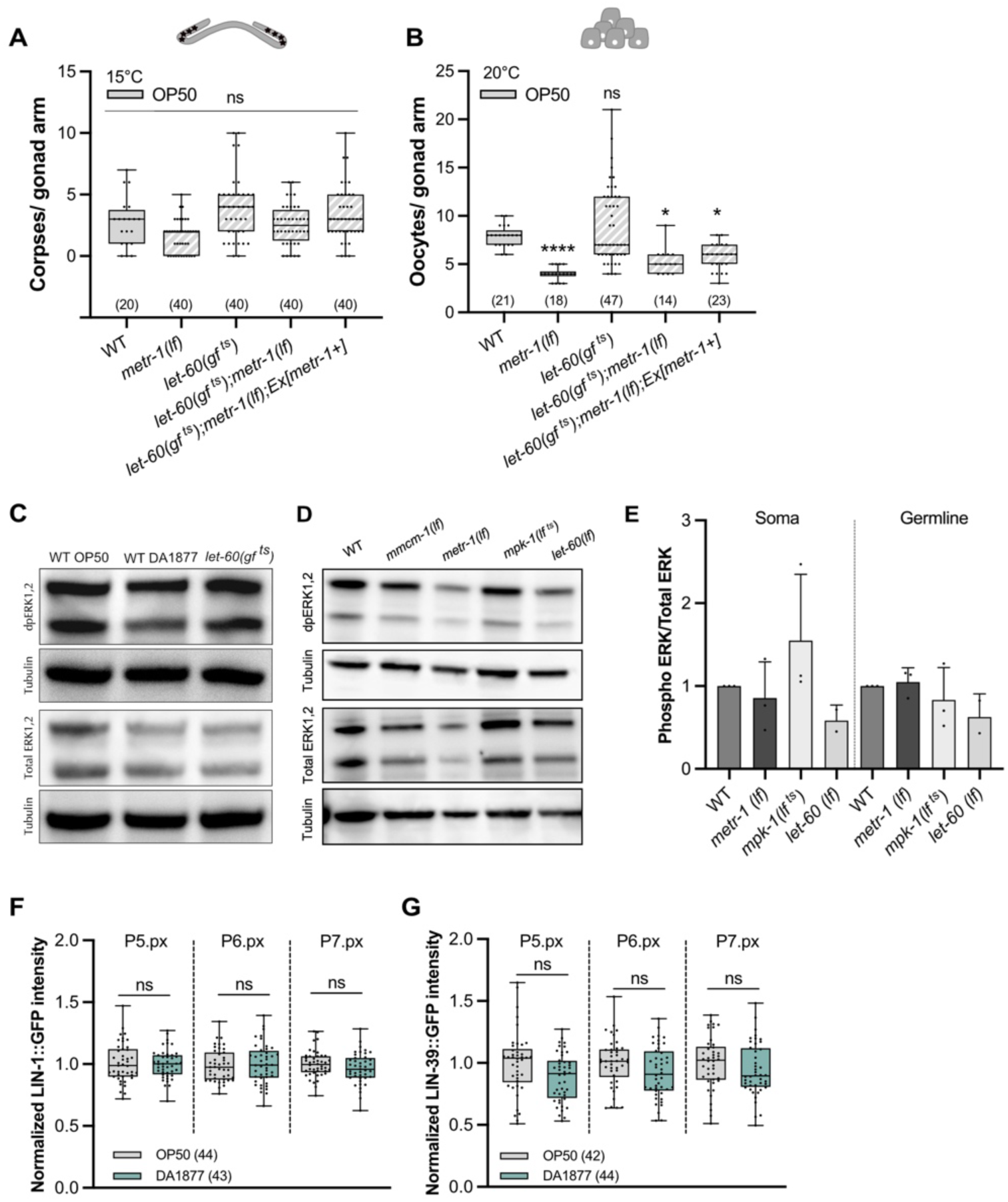
Interaction of the one-carbon metabolism and the RAS/MAPK pathway. (A-B) Number of corpses at 15°C (A) or oocytes at 20°C (B) for indicated genotype fed with OP50; dots represent individual corpses/oocytes; data represents median ± min and max from 2 independent biological replicates. (C-D) Western blot for phosphorylated and total ERK for indicated genotypes. (E) Quantification of phosphorylated and total ERK for the somatic and germline isoforms for indicated genotypes; bars represent mean ± SD normalized to the control (WT OP50) from 3 independent biological and technical replicates. (F-G) Normalized intensity of LIN-1::GFP (F) or LIN-39::GFP (G) in 1° and 2° VPC at the two-cell Pn.px stage in the wild-type fed with OP50 (gray) or DA1877 (blue); data was normalized to the control (OP50); dots represent individual animals from 2 independent biological replicates; data represents median ± min and max. (A-G) Number of animals shown in brackets. For statistical analysis see STAR methods. *<0.05; ns, non-significant.

**Figure S5.**
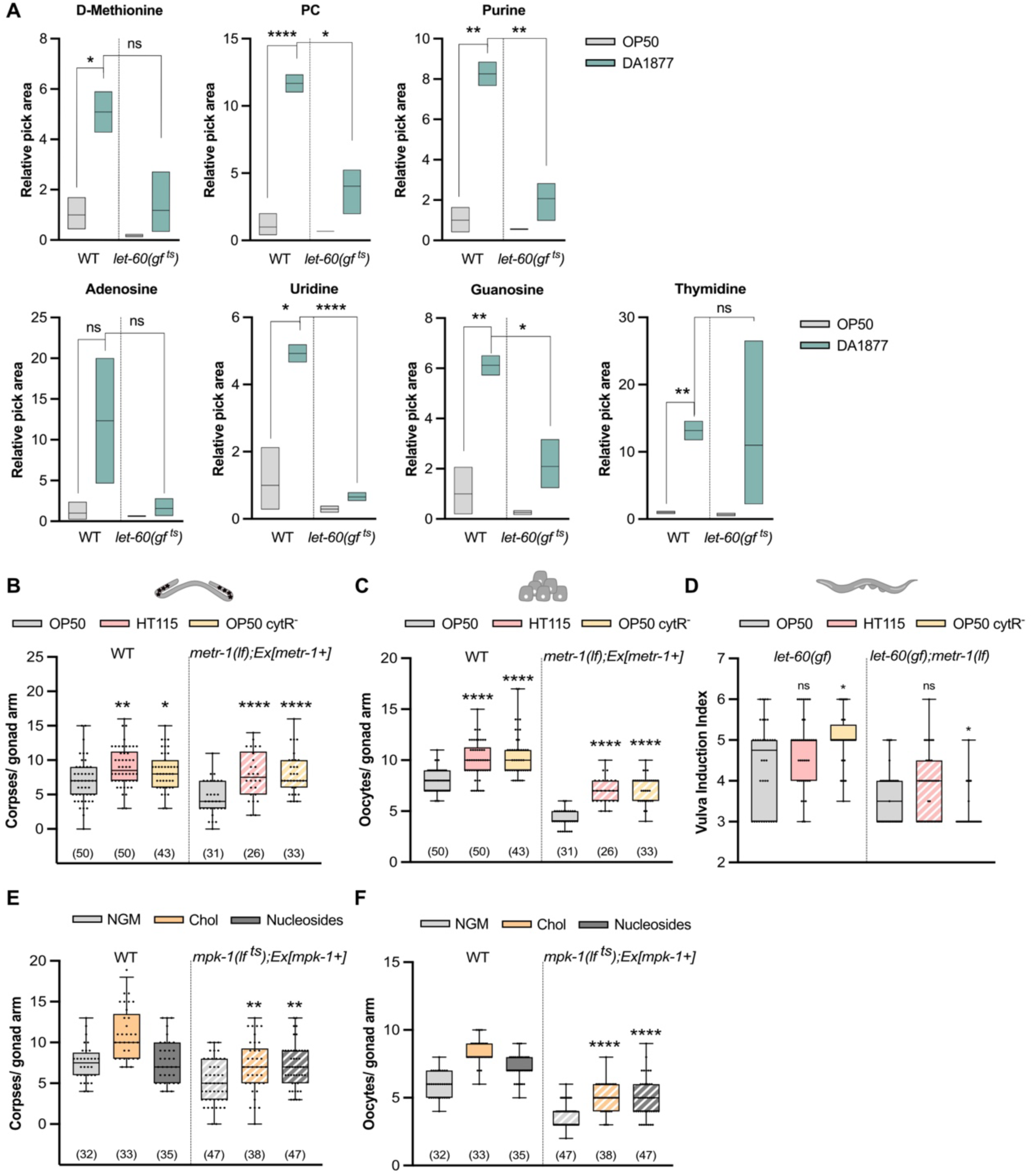
One-carbon metabolites regulate VPC and germ cell differentiation. (A) Relative levels of selected metabolites in wild-type and *let-60(gf ^ts^)* animals fed with OP50 (gray) or DA1877 (blue); *let-60(gf ^ts^)* animals were transferred to 25°C 4 hours before sample collection; peak areas were normalized to the control (wild-type OP50); bars represent mean ± min and max from 2 or 3 independent biological replicates. (B-D) Number of apoptotic corpses (B), oocytes (C) and vulval induction index (D) for indicated genotypes fed with *E. coli* OP50 (gray), *E. coli* HT115 (pink) or *E. coli* OP50 cytR^-^ (yellow). (E, F) Number of apoptotic corpses (E) and oocytes (F) for indicated genotypes fed with OP50 and supplemented with 40 mM choline (orange) or 1 mM nucleosides (dark gray). Dots represent individual corpses/oocytes or animals in (D) from 2 independent biological replicates; data represents median ± min and max; number of animals represented in brackets. For statistical analysis see STAR methods. **** <0.0001; **<0.005; *<0.05; ns, non-significant.

**Figure S6.**
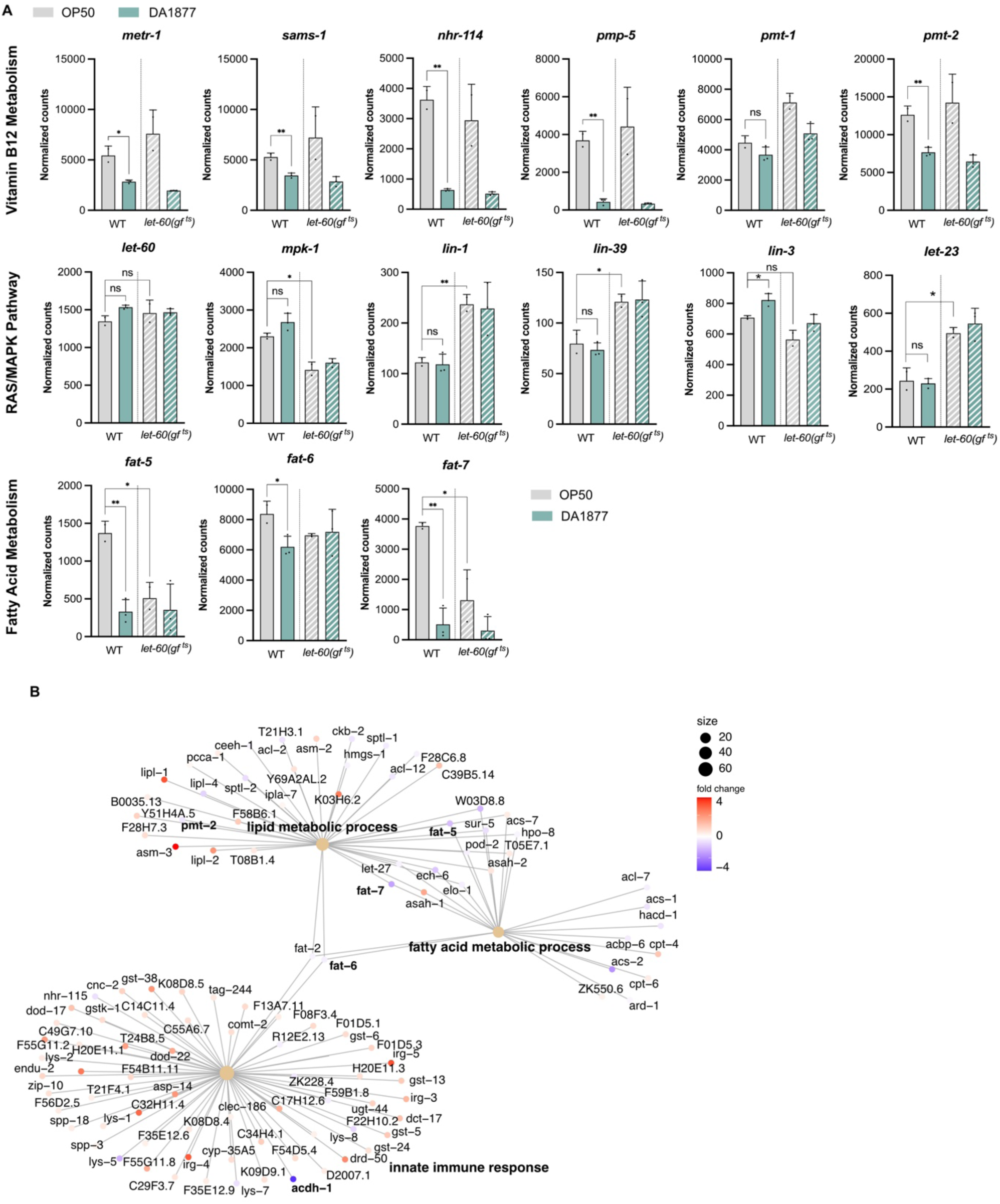
Effect of the DA1877 diet on gene expression in adults. (A) Differentially expressed genes in wild-type and *let-60(gf ^ts^)* animals fed with OP50 (gray) or DA1877 (blue); *let-60(gf ^ts^)* animals were transferred to 25°C 4 hours before sample collection; bars represent mean ± SD. (B) Pathway analysis of genes most strongly affected by the DA1877 diet in wild-type worms; circle sizes represent normalized counts and colors the fold change (see legend). For statistical analysis, see STAR methods. **<0.005; **<0.005; ns, non-significant.

**Figure S7.**
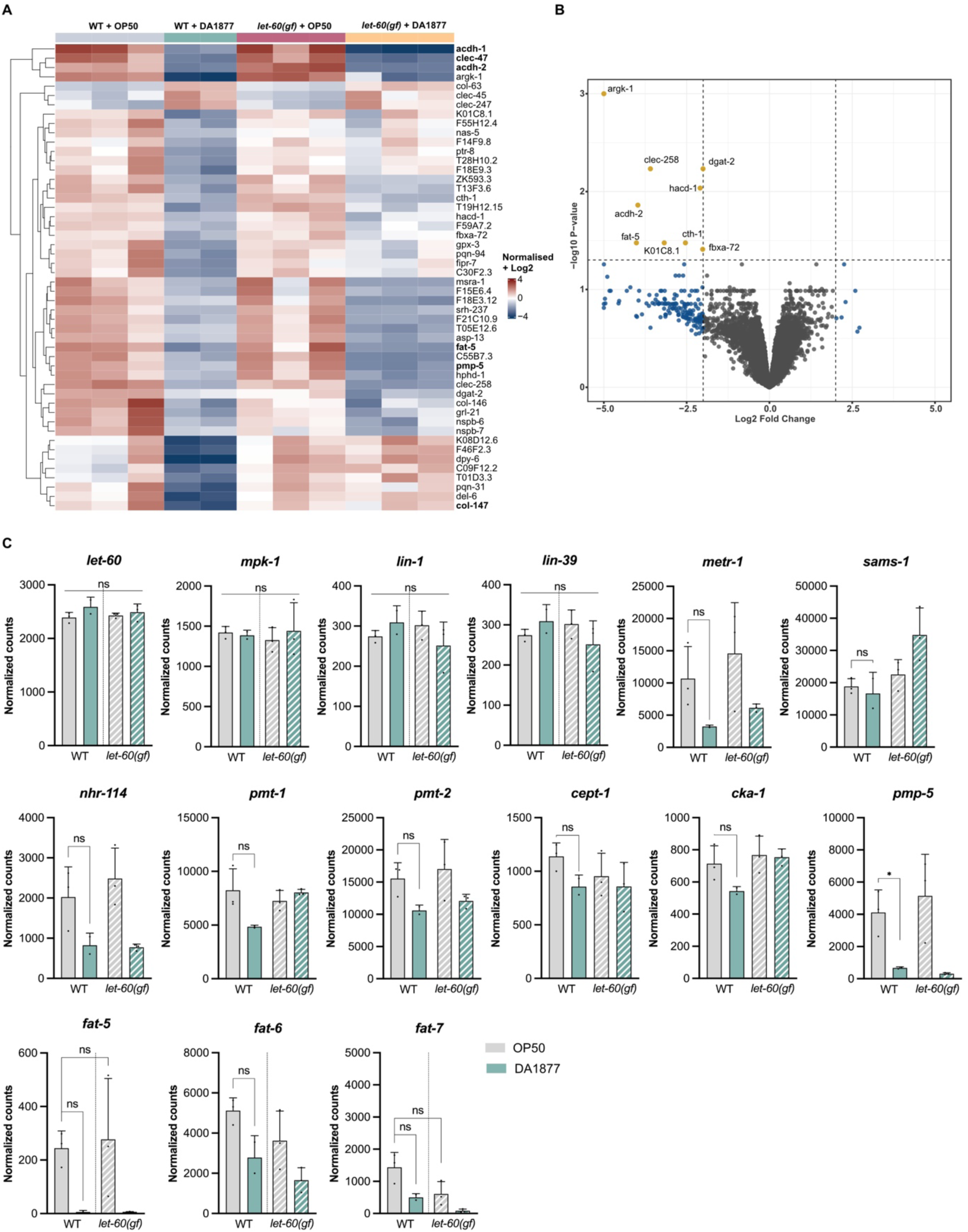
Effect of the DA1877 diet on gene expression in L3 larvae. (A) Heat map of the top 50 differentially expressed genes for indicated genotypes. (B) Differentially expressed genes in L3 larvae fed with OP50 or DA1877; p-value calculated with FDR values and threshold for p-value 0.05 are shown. (C) Differentially expressed selected genes in wild-type and *let-60(gf)* animals fed with OP50 (gray) or DA1877 (blue); bars represent mean ± SD. For statistical analysis, see STAR methods. *<0.05; ns, non-significant.

**Figure S8.**
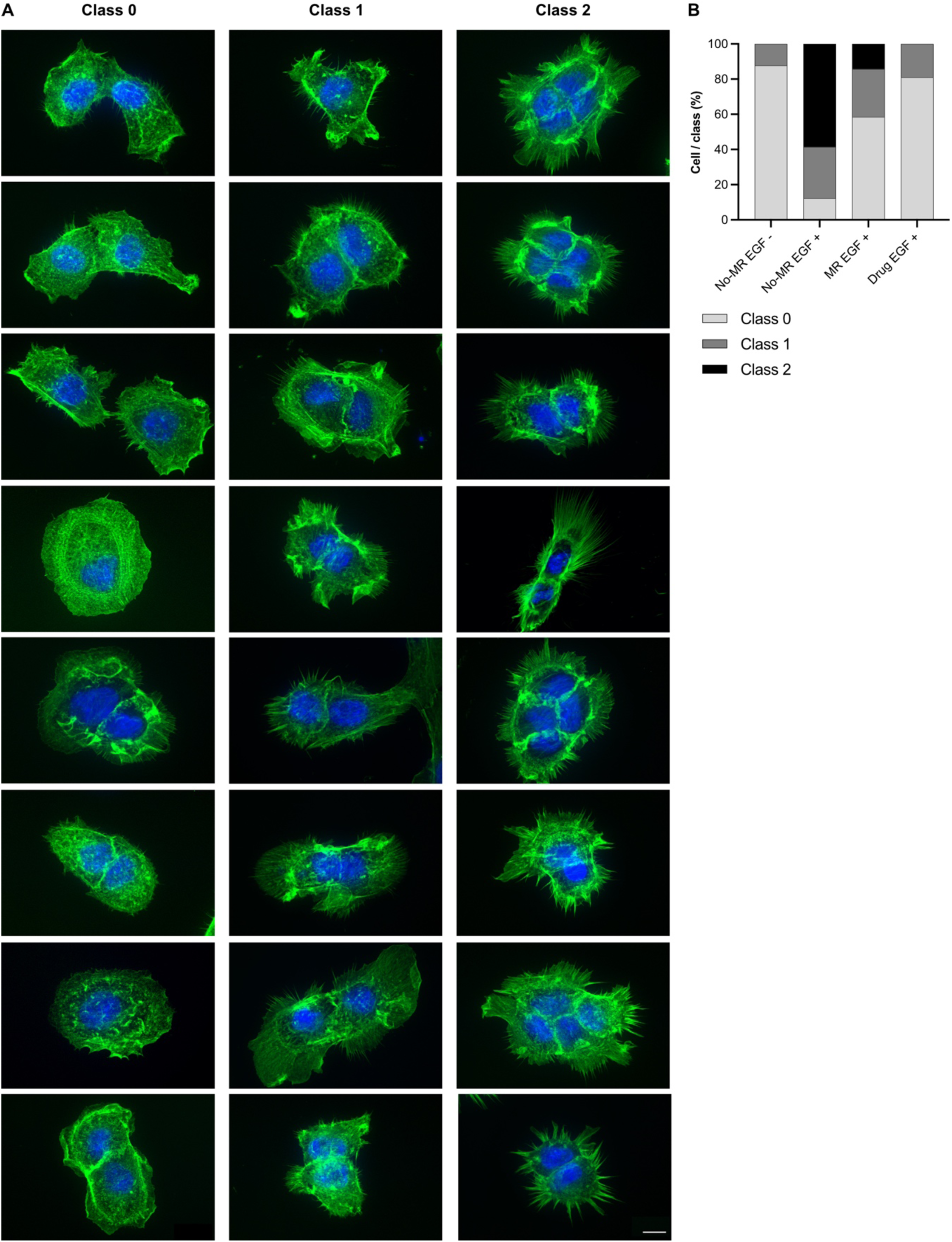
Methionine dependency of RAS/MAPK-induced phenotypes in mammalian cells. (A) Examples of A431 cells stained with DAPI (blue) and phalloidin (green) after EGF stimulation; cells were serum starved for 16 hours and stimulated for 10 min with 100 ng/ml human EGF before fixation; cells were categorized into three qualitative groups: 0 for cells without filopodia, 1 for cells with short filopodia not covering the whole cell surface and 2 for cells with long filopodia covering most of the cell surface; the panel shows examples for each class; scale bar: 10 µm. (B) Percentage of cells in each class with and without MR, with and without EGF-stimulation or MEK inhibitors as indicated on the x-axis.

## Key resources table

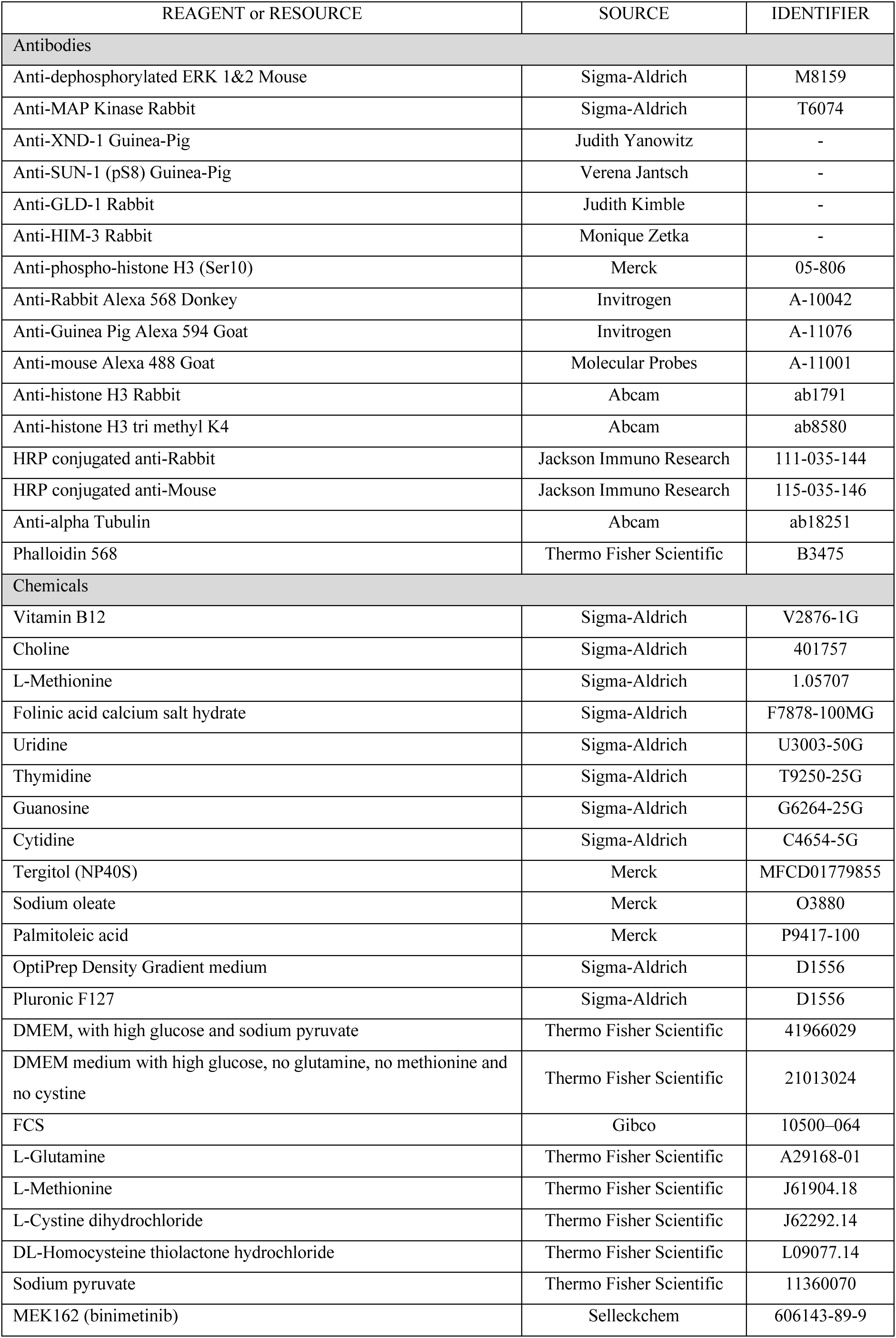
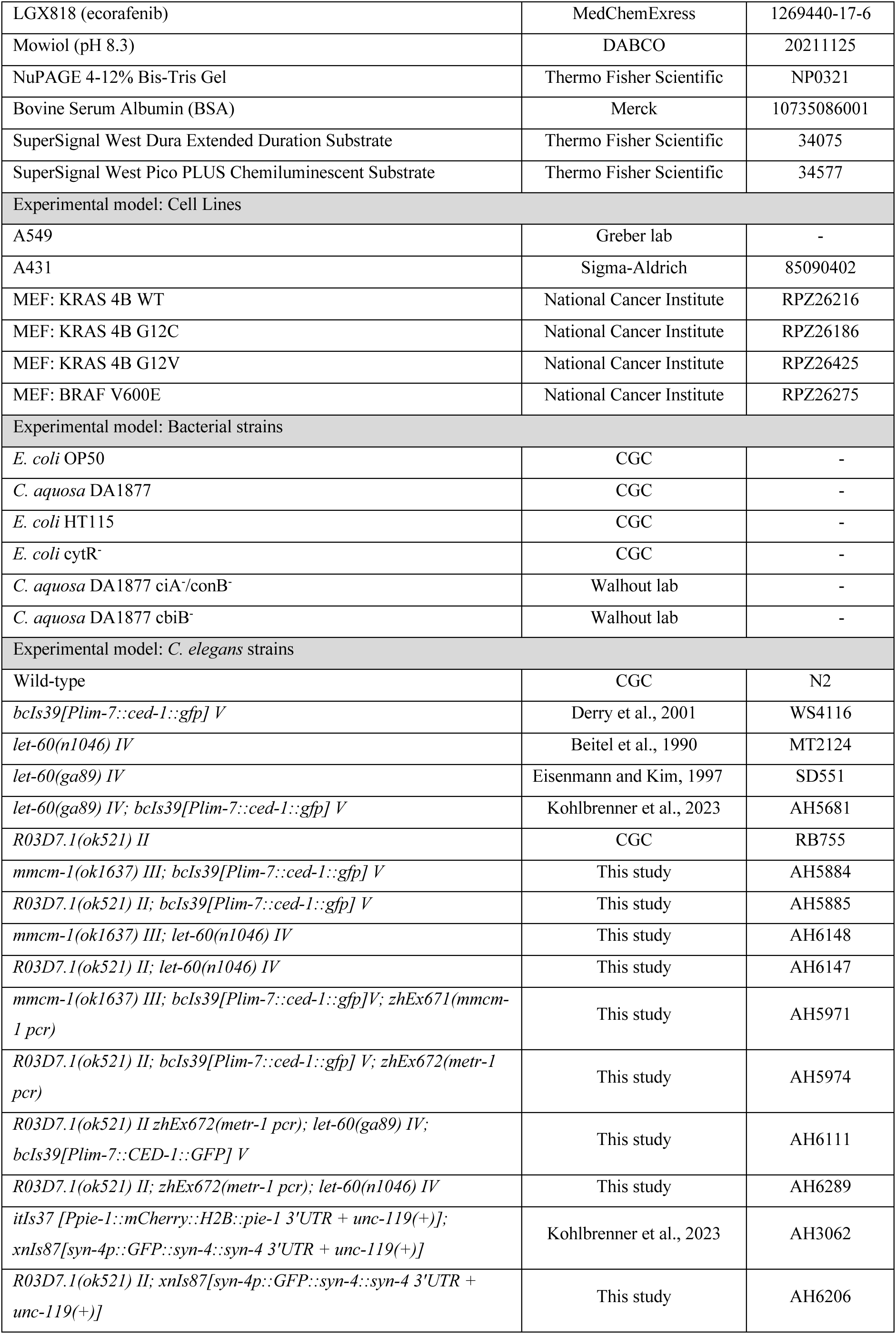
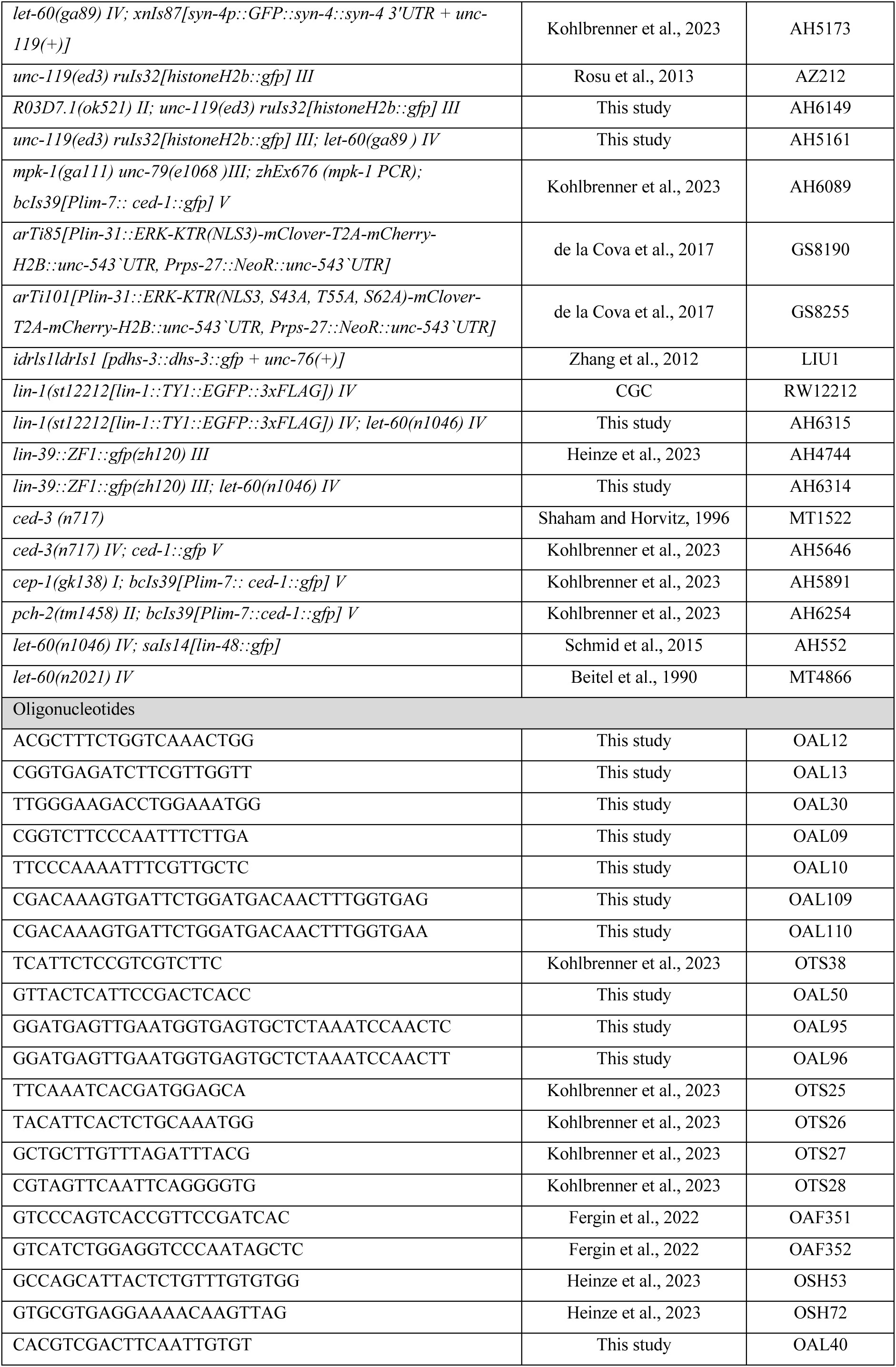
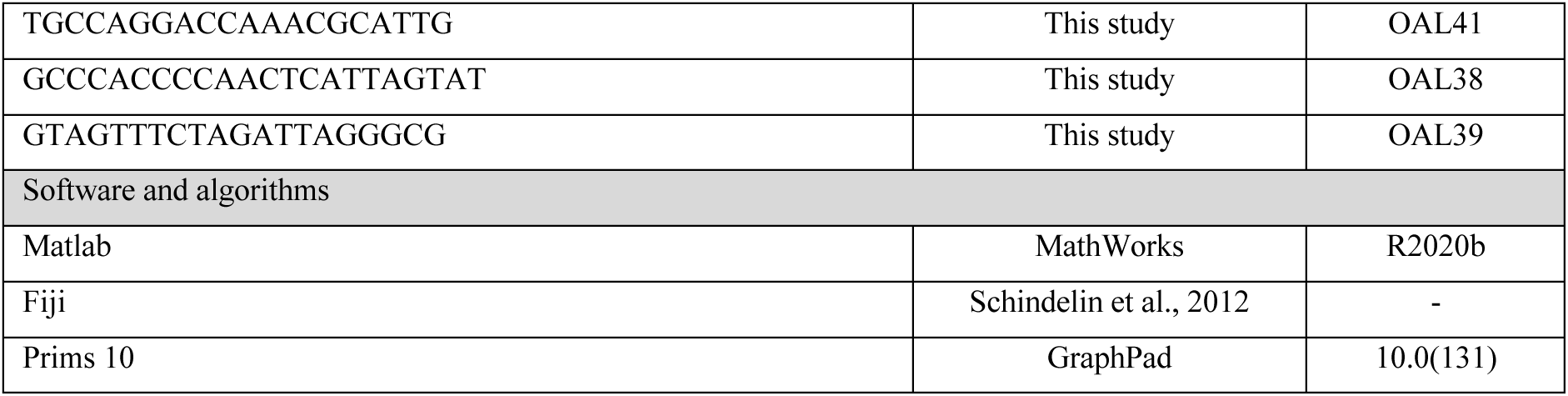

## Notes

### Competing Interest Statement

The authors have declared no competing interest.

## References

Anthony, M.L., Zhao, M., and Brindle, K.M. (1999). Inhibition of phosphatidylcholine biosynthesis following induction of apoptosis in HL-60 cells. J. Biol. Chem. 274, 19686–19692.

Arur, S., Ohmachi, M., Berkseth, M., Nayak, S., Hansen, D., Zarkower, D., and Schedl, T. (2011). MPK-1 ERK controls membrane organization in C. elegans oogenesis via a sex-determination module. Dev. Cell 20, 677– 688.

Beitel, G.J., Clark, S.G., and Horvitz, H.R. (1990). Caenorhabditis elegans ras gene let-60 acts as a switch in the pathway of vulval induction. Nature 348, 503–509.

Berger, S., Lattmann, E., Aegerter-Wilmsen, T., Hengartner, M., Hajnal, A., Demello, A., and Casadevall I Solvas, X. (2018). Long-term: C. elegans immobilization enables high-resolution developmental studies in vivo. Lab Chip 18, 1359–1368.

Bito, T., and Watanabe, F. (2016). Biochemistry, function, and deficiency of vitamin B12 in Caenorhabditis elegans. Exp. Biol. Med. 241, 1663–1668.

Booher, K., Lin, D.-W., Borrego, S.L., and Kaiser, P. (2012). Downregulation of Cdc6 and pre-replication complexes in response to methionine stress in breast cancer cells. Cell Cycle 11, 4414–4423.

Bose, S., Allen, A.E., and Locasale, J.W. (2020). The Molecular Link from Diet to Cancer Cell Metabolism. Mol. Cell 78, 1034–1044.

Brendza, K.M., Haakenson, W., Cahoon, R.E., Hicks, L.M., Palavalli, L.H., Chiapelli, B.J., McLaird, M., McCarter, J.P., Williams, D.J., Hresko, M.C., et al. (2007). Phosphoethanolamine N-methyltransferase (PMT-1) catalyses the first reaction of a new pathway for phosphocholine biosynthesis in Caenorhabditis elegans. Biochem. J. 404, 439–448.

Brenner, S. (1974). The genetics of Caenorhabditis elegans. Genetics 77, 71–94.

Brock, T.J., Browse, J., and Watts, J.L. (2007). Fatty acid desaturation and the regulation of adiposity in Caenorhabditis elegans. Genetics 176, 865–875.

Cavuoto, P., and Fenech, M.F. (2012). A review of methionine dependency and the role of methionine restriction in cancer growth control and life-span extension. Cancer Treat. Rev. 38, 726–736.

Cha, D.S., Datla, U.S., Hollis, S.E., Kimble, J., and Lee, M.H. (2012). The Ras-ERK MAPK regulatory network controls dedifferentiation in Caenorhabditis elegans germline. Biochim. Biophys. Acta - Mol. Cell Res. 1823, 1847–1855.

Chaudhari, S.N., Mukherjee, M., Vagasi, A.S., Bi, G., Rahman, M.M., Nguyen, C.Q., Paul, L., Selhub, J., and Kipreos, E.T. (2016). Bacterial Folates Provide an Exogenous Signal for C. elegans Germline Stem Cell Proliferation. Dev. Cell 38, 33–46.

Chello, P.L., and Bertino, J.R. (1973). Dependence of 5-Methyltetrahydrofolate Utilization by L5178Y Murine Leukemia Cells in Vitro on the Presence of Hydroxycobalamin and Transcobalamin II. Cancer Res. 33, 1989–1904.

Chi, C., Ronai, D., Than, M.T., Walker, C.J., Sewell, A.K., and Han, M. (2016). Nucleotide levels regulate germline proliferation through modulating GLP-1/Notch signaling in C. elegans. Genes Dev. 30, 307–320.

Church, D.L., Guan, K.L., and Lambie, E.J. (1995). Three genes of the MAP kinase cascade, mek-2, mpk1/sur-1 and let-60 ras, are required for meiotic cell cycle progression in Caenorhabditis elegans. Development 121, 2525–2535.

Crook, M., Upadhyay, A., Ido, L.J., and Hanna-Rose, W. (2016). Epidermal growth factor receptor cell survival signaling requires phosphatidylcholine biosynthesis. G3 Genes, Genomes, Genet. 6, 3533–3540.

Cui, Z., and Houweling, M. (2002). Phosphatidylcholine and cell death. Biochim. Biophys. Acta - Mol. Cell Biol. Lipids 1585, 87–96.

Dai, Y., Tang, H., and Pang, S. (2021). The Crucial Roles of Phospholipids in Aging and Lifespan Regulation. Front. Physiol. 12, 1–7.

Dai, Z., Mentch, S.J., Gao, X., Nichenametla, S.N., and Locasale, J.W. (2018). Methionine metabolism influences genomic architecture and gene expression through H3K4me3 peak width. Nat. Commun. 2018 91 9, 1–12.

Das, D., Chen, S.Y., and Arur, S. (2020). ERK phosphorylates chromosomal axis component HORMA domain protein HTP-1 to regulate oocyte numbers. Sci. Adv. 6.

Dave, J., Beitel, G.J., Scott, G.C., Horvitz, H.R., and Kornfeld, K. (1998). Gain-of-function mutations in the Caenorhabditis elegans lin-1 ETS gene identify a c-terminal regulatory domain phosphorylated by ERK MAP Kinase. Nature 388, 539–547.

Deline, M.L., Vrablik, T.L., and Watts, J.L. (2013). Dietary supplementation of polyunsaturated fatty acids in Caenorhabditis elegans. J. Vis. Exp. 1–7.

Deminice, R., de Castro, G.S.F., Francisco, L.V., da Silva, L.E.C.M., Cardoso, J.F.R., Frajacomo, F.T.T., Teodoro, B.G., dos Reis Silveira, L., and Jordao, A.A. (2015). Creatine supplementation prevents fatty liver in rats fed choline-deficient diet: A burden of one-carbon and fatty acid metabolism. J. Nutr. Biochem. 26, 391–397.

Deng, X., Yin, X., Allan, R., Lu, D.D., Maurer, C.W., Haimovitz-Friedman, A., Fuks, Z., Shaham, S., and Kolesnick, R. (2008). Ceramide biogenesis is required for radiation-induced apoptosis in the germ line of C. elegans. Science (80-.). 322, 110–115.

Derry, W.B., Putzke, A.P., and Rothman, J.H. (2001). Caenorhabditis elegans p53: Role in apoptosis, meiosis, and stress resistance. Science (80-.). 294, 591–595.

Ding, W., Smulan, L.J., Hou, N.S., Taubert, S., Watts, J.L., and Walker, A.K. (2015). S-adenosylmethionine levels govern innate immunity through distinct methylation-dependent pathways. Cell Metab. 22, 633–645.

Ding, W., Higgins, D.P., Yadav, D.K., Godbole, A.A., Pukkila-Worley, R., and Walker, A.K. (2018). Stress-responsive and metabolic gene regulation are altered in low S-adenosylmethionine. PLoS Genet. 14, 1–26.

Diot, C., García-González, A.P., Vieira, A.F., Walker, M., Honeywell, M., Doyle, H., Ponomarova, O., Rivera, Y., Na, H., Zhang, H., et al. (2022). Bacterial diet modulates tamoxifen-induced death via host fatty acid metabolism. Nat. Commun. 13.

Drosten, M., Dhawahir, A., Sum, E.Y.M., Urosevic, J., Lechuga, C.G., Esteban, L.M., Castellano, E., Guerra, C., Santos, E., and Barbacid, M. (2010). Genetic analysis of Ras signalling pathways in cell proliferation, migration and survival. EMBO J. 29, 1091–1104.

Ducker, G.S., and Rabinowitz, J.D. (2017). One-Carbon Metabolism in Health and Disease. Cell Metab. 25, 27–42.

Eisenmann, D.M., and Kim, S.K. (1997). Mechanism of Activation of the Caenorhabditis elegans rm Homologue let-60 by a Novel, Temperature-Sensitive, Gain-of-Function Mutation. Genetics 565, 553–565.

Fergin, A., Boesch, G., Greter, N.R., Berger, S., and Hajnal, A. (2022). Tissue-specific inhibition of protein sumoylation uncovers diverse SUMO functions during C. elegans vulval development. PLoS Genet. 18, 1–22.

Fisher, K., Southall, S.M., Wilson, J.R., and Poulin, G.B. (2010). Methylation and demethylation activities of a C. elegans MLL-like complex attenuate RAS signalling. Dev. Biol. 341, 142–153.

Froese, D.S., Fowler, B., and Baumgartner, M.R. (2019). Vitamin B12, folate, and the methionine remethylation cycle—biochemistry, pathways, and regulation. J. Inherit. Metab. Dis. 42, 673–685.

Frøkjaer-Jensen, C., Davis, M.W., Hopkins, C.E., Newman, B., Thummel, J.M., Olesen, S.-P., Grunnet, M., and Jorgensen, E.M. (2008). Single copy insertion of transgenes in C. elegans. Nat. Genet. 40.

Gartner, A., Milstein, S., Ahmed, S., Hodgkin, J., and Hengartner, M.O. (2000). A Conserved Checkpoint Pathway Mediates DNA Damage–Induced Apoptosis and Cell Cycle Arrest in C. elegans. Mol. Cell 5, 435–443.

Giese, G.E., Walker, M.D., Ponomarova, O., Zhang, H., Li, X., Minevich, G., and Walhout, A.J.M. (2020). Caenorhabditis elegans methionine / S-adenosylmethionine cycle activity is sensed and adjusted by a nuclear hormone receptor. Elife 1–25.

Glunde, K., Bhujwalla, Z.M., and Ronen, S.M. (2011). Choline metabolism in malignant transformation. Nat. Rev. Cancer 11, 835–848.

Godbole, A.A., Gopalan, S., Nguyen, T.K., Munden, A.L., Lui, D.S., Fanelli, M.J., Vo, P., Lewis, C.A., Spinelli, J.B., Fazzio, T.G., et al. (2023). S-adenosylmethionine synthases specify distinct H3K4me3 populations and gene expression patterns during heat stress. Elife 12, 1–26.

Goudeau, J., Bellemin, S., Toselli-Mollereau, E., Shamalnasab, M., Chen, Y., and Aguilaniu, H. (2011). Fatty acid desaturation links germ cell loss to longevity through NHR-80/HNF4 in C. elegans. PLoS Biol. 9.

Green, R., Allen, L.H., Bjørke-Monsen, A.L., Brito, A., Guéant, J.L., Miller, J.W., Molloy, A.M., Nexo, E., Stabler, S., Toh, B.H., et al. (2017). Vitamin B12 deficiency. Nat. Rev. Dis. Prim. 3.

Greenwald, I. (1997). Development of the Vulva. C. Elegans II.

Greer, E.L., Maures, T.J., Hauswirth, A.G., Green, E.M., Leeman, D.S., Maro, G.S., Han, S., Banko, M.R., Gozani, O., and Brunet, A. (2010). Members of the H3K4 trimethylation complex regulate lifespan in a germline-dependent manner in C. elegans. Nature 466, 383–387.

Gumienny, T.L., Lambie, E., Hartwieg, E., Horvitz, H.R., and Hengartner, M.O. (1999). Genetic control of programmed cell death in the Caenorhabditis elegans hermaphrodite germline. 1022, 1011–1022.

Guo, Y., Pan, W., Liu, S., Shen, Z., Xu, Y., and Hu, L. (2020). ERK/MAPK signalling pathway and tumorigenesis (Review). Exp. Ther. Med. 1997–2007.

Gupta, B.P., Hanna-Rose, W., and Sternberg, P.W. (2012). Morphogenesis of the vulva and the vulval-uterine connection*. Wormbook.

Haag, A., Walser, M., Henggeler, A., and Hajnal, A. (2020). The CHORD protein CHP-1 regulates EGF receptor trafficking and signaling in C. elegans and in human cells. Elife 9, 1–23.

Han, M., Aroian, R. V, and Sternberg, P.W. (1990). The let-60 Locus Controls the Switch Between Vulval and Nonvulval Cell Fates in Caenorhabditis elegans.

Hanahan, D., and Weinberg, R.A. (2011). Hallmarks of cancer: The next generation. Cell 144, 646–674.

Hidalgo, M.A., Nahuelpan, C., Manosalva, C., Jara, E., Carretta, M.D., Conejeros, I., Loaiza, A., Chihuailaf, R., and Burgos, R.A. (2011). Oleic acid induces intracellular calcium mobilization, MAPK phosphorylation, superoxide production and granule release in bovine neutrophils. Biochem. Biophys. Res. Commun. 409, 280–286.

Huelgas-Morales, G., and Greenstein, D. (2018). Control of oocyte meiotic maturation in C. elegans. Semin. Cell Dev. Biol. 84, 90–99.

Kaiser, P. (2020). Methionine dependence of cancer. Biomolecules 10, 7–9.

Kera, Y., Katoh, Y., Ohta, M., Matsumoto, M., Takano-Yamamoto, T., and Igarashi, K. (2013). Methionine adenosyltransferase II-dependent histone H3K9 methylation at the COX-2 gene locus. J. Biol. Chem. 288, 13592– 13601.

Kohlbrenner, T., Berger, S., Aegerter-Wilmsen, T., Laranjeira, A.C., DeMello, A., and Hajnal, A. (2023). Actomyosin-mediated apical constriction promotes physiological germ cell death in C. elegans.

Krueger, J.S., Keshamouni, V.G., Atanaskova, N., and Reddy, K.B. (2001). Temporal and quantitative regulation of mitogen-activated protein kinase (MAPK) modulates cell motility and invasion. Oncogene 20, 4209– 4218.

de la Cova, C., Townley, R., Regot, S., and Greenwald, I. (2017). A Real-Time Biosensor for ERK Activity Reveals Signaling Dynamics during C. elegans Cell Fate Specification. Dev. Cell 42, 542–553.e4.

Lackner, M.R., and Kim, S.K. (1998). Genetic analysis of the Caenorhabditis elegans MAP kinase gene mpk-1. Genetics 150, 103–117.

Lee, H.S., Nam, Y., Chung, Y.H., Kim, H.R., Park, E.S., Chung, S.J., Kim, J.H., Sohn, U.D., Kim, H.C., Oh, K.W., et al. (2014). Beneficial effects of phosphatidylcholine on high-fat diet-induced obesity, hyperlipidemia and fatty liver in mice. Life Sci. 118, 7–14.

Lee, M.H., Ohmachi, M., Arur, S., Nayak, S., Francis, R., Church, D., Lambie, E., and Schedl, T. (2007). Multiple functions and dynamic activation of MPK-1 extracellular signal-regulated kinase signaling in Caenorhabditis elegans germline development. Genetics 177, 2039–2062.

Lemieux, G.A., and Ashrafi, K. (2016). Investigating Connections between Metabolism, Longevity, and Behavior in Caenorhabditis elegans. Trends Endocrinol. Metab. 27, 586–596.

Li, T., and Kelly, W.G. (2011). A role for Set1/MLL-related components in epigenetic regulation of the Caenorhabditis elegans germ line. PLoS Genet. 7.

Li, S., Swanson, S.K., Gogol, M., Florens, L., Washburn, M.P., Workman, J.L., and Suganuma, T. (2015). Serine and SAM Responsive Complex SESAME Regulates Histone Modification Crosstalk by Sensing Cellular Metabolism. Mol. Cell 60, 408–421.

Li, Y., Na, K., Lee, H.J., Lee, E.Y., and Paik, Y.K. (2011). Contribution of sams-1 and pmt-1 to lipid homeostasis in adult Caenorhabditis elegans. J. Biochem. 149, 529–538.

Liu, F., Yang, X., Geng, M., and Huang, M. (2018). Targeting ERK, an Achilles’ Heel of the MAPK pathway, in cancer therapy. Acta Pharm. Sin. B 8, 552–562.

Lochnit, G., and Geyer, R. (2003). Evidence for the presence of the Kennedy and Bremer-Greenberg pathways in Caenorhabditis elegans. Acta Biochim. Pol. 50, 1239–1243.

MacNeil, L.T., Watson, E., Arda, H.E., Zhu, L.J., and Walhout, A.J.M. (2013). Diet-induced developmental acceleration independent of TOR and insulin in C. elegans. Cell 153, 240–252.

Mello1, C.C., Kramer2’, J.M., Stinchcomb4, D., and Ambros, V. (1991). Efficient gene transfer in C.elegans: extrachromosomal maintenance and integration of transforming sequences. EMBO J. 10, 3959–3970.

Mentch, S.J., Mehrmohamadi, M., Huang, L., Liu, X., Gupta, D., Mattocks, D., Gómez Padilla, P., Ables, G., Bamman, M.M., Thalacker-Mercer, A.E., et al. (2015). Histone Methylation Dynamics and Gene Regulation Occur through the Sensing of One-Carbon Metabolism. Cell Metab. 22, 861–873.

Newman, A.C., and Maddocks, O.D.K. (2017). One-carbon metabolism in cancer. Br. J. Cancer 116, 1499– 1504.

Niculescu, M.D., Craciunescu, C.N., and Zeisel, S.H. (2006). Dietary choline deficiency alters global and gene-specific DNA methylation in the developing hippocampus of mouse fetal brains. FASEB J. • Res. Commun.

Özdemir, I., and Steiner, F.A. (2022). Transmission of chromatin states across generations in C. elegans. Semin. Cell Dev. Biol. 127, 133–141.

Pazdernik, N., and Sched, T. (2013). Introduction to Germ Cell Development in C. elegans. Adv. Exp. Med. Biol. 757, 249–276.

Perez, M.A., and Watts, J.L. (2021). Worms, fat, and death: Caenorhabditis elegans lipid metabolites regulate cell death. Metabolites 11, 1–18.

Perez, M.A., Magtanong, L., Dixon, S.J., and Watts, J.L. (2020). Dietary Lipids Induce Ferroptosis in Caenorhabditiselegans and Human Cancer Cells. Dev. Cell 54, 447–454.e4.

Qin, S., Wang, Y., Li, L., Liu, J., Xiao, C., Duan, D., Hao, W., Qin, C., Chen, J., Yao, L., et al. (2022). Early-life vitamin B12 orchestrates lipid peroxidation to ensure reproductive success via SBP-1/SREBP1 in Caenorhabditis elegans. Cell Rep. 40, 111381.

Robinson-Thiewes, S., Dufour, B., Martel, P.O., Lechasseur, X., Brou, A.A.D., Roy, V., Chen, Y., Kimble, J., and Narbonne, P. (2021). Non-autonomous regulation of germline stem cell proliferation by somatic MPK-1/MAPK activity in C. elegans. Cell Rep. 35, 109162.

Saito, R. de F., Andrade, L.N. de S., Bustos, S.O., and Chammas, R. (2022). Phosphatidylcholine-Derived Lipid Mediators: The Crosstalk Between Cancer Cells and Immune Cells. Front. Immunol. 13, 1–24.

Santa-María, C., López-Enríquez, S., Montserrat-de la Paz, S., Geniz, I., Reyes-Quiroz, M.E., Moreno, M., Palomares, F., Sobrino, F., and Alba, G. (2023). Update on Anti-Inflammatory Molecular Mechanisms Induced by Oleic Acid. Nutrients 15, 1–16.

Schindelin, J., Arganda-Carreras, I., Frise, E., Kaynig, V., Longair, M., Pietzsch, T., Preibisch, S., Rueden, C., Saalfeld, S., Schmid, B., et al. (2012). Fiji: An open-source platform for biological-image analysis. Nat. Methods 9, 676–682.

Schindler, A.J., and Sherwood, D.R. (2013). Morphogenesis of the Caenorhabditis elegans vulva. Wiley Interdiscip. Rev. Dev. Biol. 2, 75–95.

Schmid, T., Snoek, L.B., Fröhli, E., van der Bent, M.L., Kammenga, J., and Hajnal, A. (2015). Systemic Regulation of RAS/MAPK Signaling by the Serotonin Metabolite 5-HIAA. PLoS Genet. 11, 1–16.

Schubbert, S., Shannon, K., and Bollag, G. (2007). Hyperactive Ras in developmental disorders and cancer. Nat. Rev. Cancer 7, 295–308.

Serefidou, M., Venkatasubramani, A.V., and Imhof, A. (2019). The Impact of One Carbon Metabolism on Histone Methylation. Front. Genet. 10, 1–7.

Shaham, S., and Horvitz, H.R. (1996). Developing Caenorhabditis elegans neurons may contain both cell-death protective and killer activities. Genes Dev. 10, 578–591.

Shyh-Chang, N., Locasale, J.W., Lyssiotis, C.A., Zheng, Y., Teo, R.Y., Ratanasirintrawoot, S., Zhang, J., Onder, T., Unternaehrer, J.J., Zhu, H., et al. (2013). Influence of threonine metabolism on S-adenosylmethionine and histone methylation. Science (80-.). 339, 222–226.

Spiri, S., Berger, S., Mereu, L., DeMello, A., and Hajnal, A. (2022). Reciprocal EGFR signaling in the anchor cell ensures precise inter-organ connection during Caenorhabditis elegans vulval morphogenesis. Development 149.

Sternberg, P.W., and Han, M. (1998). Genetics of RAS signaling in C. elegans. Trends Genet. 9525, 466–472.

Sternberg, P.W., and Horvitz, H.R. (1986). Pattern formation during vulval development in C. elegans. Cell 44, 761–772.

Sugimura, T., Birnbaum, S.M., Winitz, M., and Greenstein, J.P. (1959). Quantitative nutritional studies with water-soluble, chemically defined diets. VIII. The forced feeding of diets each lacking in one essential amino acid. Arch. Biochem. Biophys. 81, 448–455.

Sundaram, M. V (2013). Canonical RTK-Ras-ERK signaling and related alternative pathways *. Wormbook.

Van de Vijver, M.J., Kumar, R., and Mendelsohn, J. (1991). Ligand-induced activation of A431 cell epidermal growth factor receptors occurs primarily by an autocrine pathway that acts upon receptors on the surface rather than intracellularly. J. Biol. Chem. 266, 7503–7508.

Walker, A.K., Boag, P.R., and Blackwell, T.K. (2007). Transcription reactivation steps stimulated by oocyte maturation in C. elegans. Dev. Biol. 304, 382–393.

Walker, A.K., Jacobs, R.L., Watts, J.L., Rottiers, V., Jiang, K., Finnegan, D.M., Shioda, T., Hansen, M., Yang, F., Niebergall, L.J., et al. (2011). A conserved SREBP-1/phosphatidylcholine feedback circuit regulates lipogenesis in metazoans. Cell 147, 840–852.

Wan, Q.L., Meng, X., Fu, X., Chen, B., Yang, J., Yang, H., and Zhou, Q. (2019). Intermediate metabolites of the pyrimidine metabolism pathway extend the lifespan of C. elegans through regulating reproductive signals. Aging (Albany. NY). 11, 3993–4010.

Wan, Q.L., Meng, X., Wang, C., Dai, W., Luo, Z., Yin, Z., Ju, Z., Fu, X., Yang, J., Ye, Q., et al. (2022). Histone H3K4me3 modification is a transgenerational epigenetic signal for lipid metabolism in Caenorhabditis elegans. Nat. Commun. 13, 1–14.

Wang, X., and Yang, C. (2016). Programmed cell death and clearance of cell corpses in Caenorhabditis elegans. Cell. Mol. Life Sci. 73, 2221–2236.

Wasmuth, J., Schmid, R., Hedley, A., and Blaxter, M. (2008). On the extent and origins of genic novelty in the phylum nematoda. PLoS Negl. Trop. Dis. 2.

Watson, E., MacNeil, L.T., Arda, H.E., Zhu, L.J., and Walhout, A.J.M. (2013). Integration of metabolic and gene regulatory networks modulates the C. elegans dietary response. Cell 153, 253–266.

Watson, E., Macneil, L.T., Ritter, A.D., Yilmaz, L.S., Rosebrock, A.P., and Caudy, A.A. (2014). Interspecies Systems Biology Uncovers Metabolites Affecting C. elegans Gene Expression and Life History Traits. Cell 156, 759–770.

Watson, E., Olin-Sandoval, V., Louisse, T., Walhout, A.J., Troyanskaya, O.G., Li, C.-H., Watson, E., Holdorf, A.D., Yao, V., Ralser, M., et al. (2016). Metabolic network rewiring of propionate flux compensates vitamin B12 deficiency in C. elegans. Elife 5, 1–21.

Watts, J.L. (2009). Fat synthesis and adiposity regulation in Caenorhabditis elegans. Trends Endocrinol. Metab. 20, 58–65.

Watts, J.L., and Ristow, M. (2017). Lipid and carbohydrate metabolism in Caenorhabditis elegans. Genetics 207, 413–446.

Xiao, Y., Bedet, C., Robert, V.J.P., Simonet, T., Dunkelbarger, S., Rakotomalala, C., Soete, G., Korswagen, H.C., Strome, S., and Palladino, F. (2011). Caenorhabditis elegans chromatin-associated proteins SET-2 and ASH-2 are differentially required for histone H3 Lys 4 methylation in embryos and adult germ cells. Proc. Natl. Acad. Sci. U. S. A. 108, 8305–8310.

Yamasaki, M., Tachibana, H., Yamada, A., Ochi, Y., Madhyastha, H., Nishiyama, K., and Yamada, K. (2008). Oleic acid prevents apoptotic cell death induced by trans10, cis12 isomer of conjugated linoleic acid via p38 MAP kinase dependent pathway. Vitr. Cell. Dev. Biol. - Anim. 44, 290–294.

Yang, Y., Xu, G., Xu, Y., Cheng, X., Xu, S., Chen, S., and Wu, L. (2021). Ceramide mediates radiation-induced germ cell apoptosis via regulating mitochondria function and MAPK factors in Caenorhabditis elegans. Ecotoxicol. Environ. Saf. 208, 111579.

Ye, C., Sutter, B.M., Wang, Y., Kuang, Z., and Tu, B.P. (2017). A Metabolic Function for Phospholipid and Histone Methylation. Mol. Cell 66, 180–193.e8.

Yuan, Z., Feng, L., Jiang, W., Wu, P., Liu, Y., Kuang, S., Tang, L., and Zhou, X. (2021). Dietary choline deficiency aggravated the intestinal apoptosis in association with the MAPK signalling pathways of juvenile grass carp (Ctenopharyngodon idella). Aquaculture 532, 736046.

Zeisel, S.H. (2011). The supply of choline is important for fetal progenitor cells. Semin. Cell Dev. Biol. 22, 624–628.

Zhang, B., Zeng, M., Wang, Y., Li, M., Wu, Y., Xu, R., Zhang, Q., Jia, J., Huang, Y., Zheng, X., et al. (2022). Oleic acid alleviates LPS-induced acute kidney injury by restraining inflammation and oxidative stress via the Ras/MAPKs/PPAR-γ signaling pathway. Phytomedicine 94, 153818.

Zhang, P., Na, H., Liu, Z., Zhang, S., Xue, P., Chen, Y., Pu, J., Peng, G., Huang, X., Yang, F., et al. (2012). Proteomic study and marker protein identification of Caenorhabditis elegans lipid droplets. Mol. Cell. Proteomics 11, 317–328.

Zhou, Z., Hartwieg, E., and Horvitz, H.R. (2001). CED-1 is a transmembrane receptor that mediates cell corpse engulfment in C. elegans. Cell 104, 43–56.

Zhu, X., Liu, Y., Zhang, H., and Liu, P. (2018). Whole-genome RNAi screen identifies methylation-related genes influencing lipid metabolism in Caenorhabditis elegans. J. Genet. Genomics 45, 259–272.

